# A multi-plant transcriptomic atlas reveals conserved and lineage specific defense architectures in response to *Botrytis cinerea*

**DOI:** 10.64898/2026.01.14.699558

**Authors:** Ritu Singh, Anna Jo Muhich, Cloe Tom, Celine Caseys, Daniel J Kliebenstein

## Abstract

Generalist pathogens pose a challenge to plant immunity by infecting diverse hosts while harboring extensive intraspecific genetic variation. Whether evolutionary distant plant lineages rely on a shared immune strategy or deploy distinct, lineage-specific defenses when confronted by these genetically variable members of the same pathogen species remains unresolved. Here, we employed a large-scale co-transcriptomic approach to map the immune landscape of ten diverse eudicot species infected with 72 genetically distinct *Botrytis cinerea* isolates. We identified a limited core of evolutionarily conserved defense orthologs, along with a vast landscape of lineage-specific transcriptional rewiring. While the broad physiological outcome such as metabolic reprogramming, cell wall modification, and suppression of growth-associated processes was shared across hosts, the regulatory pathways governing this were largely lineage-specific. Crucially, this immune landscape is dynamically shaped by pathogen diversity. Nearly three-quarters of host transcriptional responses were isolate-dependent, with the magnitude of defense activation defined by specific host-isolate combinations rather than a universal species-level response. Even host responses to shared virulence factors, including broadly expressed pathogens phytotoxins, were lineage specific. These findings show that plant immunity to generalist pathogens is built on conserved physiological outcomes executed through rapidly evolving, lineage-specific regulatory programs. This distinct regulatory architecture creates an immune landscape heavily modulated by specific host-isolate combinations, highlighting the necessity of integrating pathogen diversity into models of plant defense evolution and resistance breeding.

**Significance Statement:** Achieving durable, broad-spectrum crop protection remains difficult because plant immunity models often rely on limited species and overlook natural genetic diversity. Effective crop protection requires understanding how defense networks operate across diverse lineages. We tested this by measuring immune responses of ten phylogenetically diverse eudicots infected with 72 genetically distinct *Botrytis cinerea* isolates. We found that plants share conserved physiological defense outcomes achieved through highly divergent, lineage-specific regulatory networks. Host responses were strongly shaped by pathogen genetic diversity, with identical isolates eliciting different transcriptional responses in different hosts. This demonstrates that plant immunity quantitatively senses pathogen variation and disease outcomes emerge from specific host-isolate combinations. These findings explain the limits of resistance transfer and inform strategies for durable disease control.

## Introduction

Plants are constantly challenged by a diverse array of attackers, from pathogens to herbivores, to a myriad of other pests. To survive, plants have evolved a complex and dynamic immune system capable of perceiving and counteracting biologically diverse attackers. The plant immune system operates across multiple layers of biotic interactions that can be synergistic (a defense against one attacker is effective against another attacker) or antagonistic (a defense against one attacker leads to susceptibility towards another attacker). To cope with these complex pressures, plants deploy defenses ranging from broad, innate immune responses effective against classes of attackers to highly specific mechanisms targeting individual pathogens or even distinct isolates. Evolutionarily, defenses effective against widespread threats are expected to be deeply conserved across many host species, while responses unique to specialized attackers are often lineage-specific adaptations (Bednarek et al., 2009; Clay et al., 2009; Fan et al., 2011). Understanding how this balance between conservation and diversification shapes plant immunity remains a fundamental question in evolutionary biology.

A foundation of the plant immune system is the ability to perceive signals linked to external attackers. Plant cells detect conserved molecular patterns such as microbe-, pathogen-, and damage-associated molecular patterns (MAMPs, PAMPs, DAMPs) through cell surface or cytoplasmic receptors (Jones and Dangl, 2006). This perception rapidly activates early immune responses such as ion fluxes, reactive oxygen species bursts, mitogen-activated protein kinase signaling, and callose deposition in cell walls (Windram and Denby, 2015; De Lorenzo et al., 2018; Wang et al., 2022). These early events constitute a broadly effective innate immune layer that is shared across most plants but also exhibits lineage-specific modifications. For instance, the successful functional transfer of a Barley *NLR* receptor into Arabidopsis, two species separated by ∼200 million years of evolution, demonstrates the deep conservation of underlying immune machinery (Maekawa et al., 2012). Conversely, the elongation factor-tu receptor (EFR) which recognizes the bacterial protein EF-Tu, is restricted to the Brassicaceae (Boller and Felix, 2009) illustrating how the system diversifies across lineages.

Following perception, immune signals are transmitted through intracellular signaling networks involving protein kinases, transcription factors (TFs), and phytohormone-mediated pathways. Phytohormones such as salicylic acid (SA), jasmonic acid (JA), and ethylene (ET) form central regulatory hubs to coordinate local and systemic defense responses (Pieterse et al., 2012). Although these signaling components exist across vascular plants to bryophytes (Anterola et al., 2009; P.K.G.S.S. et al., 2009; Neumann et al., 2012; Pieterse et al., 2012; PONCE DE LEÓN et al., 2012; Ponce de León and Montesano, 2013), their specific activation and crosstalk differ substantially, likely reflecting host plant tuning the distinct pathogen lifestyles and ecological contexts in which the host exists (Delaney et al., 1994; Nawrath and Métraux, 1999; Glazebrook, 2005; Windram and Denby, 2015; Hickman et al., 2019; Zander et al., 2020). For example, while the COI1-JAZ jasmonate receptor module is conserved across land plants, ligand specificity has shifted during evolution. The liverwort *Marchantia polymorpha* recognizes dn-OPDA, a precursor molecule produced early in the jasmonate biosynthetic pathway, whereas angiosperms respond to the amino acid-conjugated form jasmonoyl isoleucine (JA-Ile) (Han, 2016; Monte et al., 2018). Such flexibility allows plants to change their immune responses, leading to lineage-specific rewiring of signaling networks while maintaining a conserved core architecture.

Ultimately, immune signaling cascades coordinate the defensive outputs that directly determine infection outcomes. Some outputs such as cell wall reinforcement through lignin and callose deposition are widely conserved across plant species and function to restrict pathogen ingress. In contrast, chemical defenses are often lineage-specific, reflecting the evolutionary diversification of specialized metabolism from order to family to genera to individual species. For example, glucosinolates in Brassicaceae, camalexin in Arabidopsidae, isoflavonoids in Fabales, and cucurbitacins in Cucurbitaceae represent unique evolutionary solutions to pathogen pressure (Jeandet et al., 2014; Aeri et al., 2015). Despite extensive knowledge of individual immune components, the relative fraction of these defense components that are broadly conserved across plant families versus those that are unique to specific lineages remain elusive.

Together, the variation across plant species for the different immune layers from perception and signaling to outputs shows that the plant immune system is a mosaic of conserved and lineage-specific components. The fact that all these mechanisms show differential levels of conservation both within and between levels further complicates the ability to translate these mechanisms across plants. This raises a central question: when confronted with the same pathogen, do plant lineages, separated by millions of years of evolution, rely on a shared defense network, or has each plant species evolved distinct lineage specific defense networks against the same pathogen?

Asking this question requires measuring host responses to a pathogen that can infect diverse plant species. The necrotrophic fungus *Botrytis cinerea* provides an ideal system for this question as it can infect more than 1,500 plant species across bryophytes, monocots, and eudicots (Elad and Fillinger, 2016; Singh et al., 2024). Rather than intimately co-evolving with specific host plants, *B. cinerea* follows a generalist infection strategy that combines a core virulence mechanism activated across all hosts with host-inducible plasticity, enabling it to adapt to different hosts (You et al., 2024; Singh et al., 2025). Further, most *B. cinerea* isolates infect multiple hosts, albeit with quantitative variation in virulence. This versatility is partly underpinned by extensive intraspecific genetic variation, including major-effect polymorphisms in key virulence traits such as phytotoxin production (Colmenares et al., 2002; DALMAIS et al., 2011; Zhang et al., 2017), detoxification of host defense compounds (Ferrari et al., 2003; Pedras et al., 2005; Pedras et al., 2007; Pedras et al., 2008; Rowe et al., 2010; Pedras et al., 2011; Kuroyanagi et al., 2022; Bulasag et al., 2024), and cell wall degradation (Rowe and Kliebenstein, 2007; Schumacher, 2012; Kumari et al., 2014). This genetic variation is predominantly single nucleotide variation with little evidence for widespread presence-absence variation (Krishnan et al., 2023). Complementing this sequence variation contribution to broad host range is a capacity of *B. cinerea* to shape its transcriptome response uniquely to each host plant (Singh et al., 2025). Consequently, even within a single host species, infection outcomes differ dramatically depending on the *B. cinerea* isolate, highlighting the dynamic interplay between pathogen genetic diversity and host defenses.

This combination of broad host range and high intraspecific diversity provides a unique opportunity to ask how evolutionarily divergent plant hosts integrate pathogen genetic variation into their immune responses. A previous comparative study has tried to explore the host responses to a generalist pathogen *Sclerotinia sclerotiorum* but relied on a single pathogen isolate, limiting the ability to separate host phylogenetic effects from pathogen genetic variation (Sucher et al., 2020). To provide the capacity to assess how distinct hosts integrate across genetic variation in the pathogen, we infected ten phylogenetically diverse eudicot species with a collection of 72 genetically distinct *B. cinerea* isolates that capture a significant fraction of the species’ natural variation (Rowe and Kliebenstein, 2007; Atwell et al., 2015; Corwin et al., 2016; Zhang et al., 2017; Atwell et al., 2018; Caseys et al., 2021; Singh et al., 2025). By integrating lesion phenotyping with co-transcriptome data across hundreds of unique host-pathogen combinations, we aim to: (1) test whether the immune responses of phylogenetically diverse hosts have maintained a common set of defense mechanisms or have evolved into distinct, lineage-specific strategies; (2) disentangle the general host response to *B. cinerea* as a species from the isolate-specific effects that arise from pathogen genetic variation; and (3) correlate host gene expression with the expression of *B. cinerea* virulence factors, to map the specific host response/targets and if they are conserved or lineage-specific across host species. This framework enables a mechanistic dissection of how evolutionary divergence in both host and pathogen contributes to the architecture of plant immunity.

## Results

### Host transcriptional responses to *B. cinerea* infection

To measure how host defense responses to a single pathogen vary across eudicots we performed a co-transcriptome analysis of ten host species, each inoculated with 72 genetically diverse *B. cinerea* isolates (Rowe and Kliebenstein, 2007; Atwell et al., 2015; Corwin et al., 2016; Zhang et al., 2017; Atwell et al., 2018; Caseys et al., 2021; Singh et al., 2025). Hosts were chosen to sample across asterid and rosid clades to provide comparison at different phylogenetic levels (Singh et al., 2025). All plant species had reference genomes enabling precise RNA-seq reads alignment and quantification of host gene expression (**Table S1**). The pathogen isolates largely infect all the host plants with quantitative variation in virulence creating a population of *B. cinerea* that can test how host transcriptional responses vary across a broad phylogenetic range of eudicots (Singh et al., 2025). Further, the use of multiple isolates also allowed us to distinguish host responses common to *B. cinerea* as a species versus host responses limited to a subset of pathogen genetic variants.

To capture coordinated host and pathogen transcriptional responses during early infection, inoculated leaves were sampled at 48 hours post inoculation (hpi). RNA-seq libraries were generated for all host by isolate combinations as well as mock inoculated controls, producing a high resolution co-transcriptomic dataset that simultaneously quantified host and *B. cinerea* gene expression. Across the ten host species, we detected expression for 48% to 66% of total annotated protein-coding genes (defined as CPM > 1 in ∼ 20% of samples per species), providing a comprehensive view of the transcriptional landscape. The same dataset was previously used to analyze *B. cinerea* virulence strategies across eudicots (Singh et al., 2025). In this work we focused primarily on the host transcriptomes.

Gene expression in each host was modelled using generalized linear models with negative binomial (glmmTMB). The model included terms for overall infection (infected; uninfected versus infected), isolate-specific effects nested within infection (infected/isolate), and tray and sequencing batch covariates to correct for experimental and technical variation. This framework partitioned host responses into two components: (1) infection-responsive differentially expressed genes (DEGs), defined as host genes differentially expressed between uninfected and infected samples regardless of isolate (infected, log₂FC > 1, FDR ≤ 0.05), and (2) isolate-responsive DEGs, representing host genes whose expression varied significantly depending on the specific pathogen isolates (infected/isolate, FDR ≤ 0.05).

Across the ten hosts, the number of infection-responsive DEGs varied widely, ranging from 413 in pepper to 9,291 in lettuce (median of 28.3% ± 15.4% of expressed genes) (**Figure 1A; Table S2**). In contrast, the number of isolate-responsive host genes was more consistent across hosts, ranging from 7,345 in cucumber and 24,379 in sunflower, with most expressed genes showing significant isolate-dependent effects (median of 73.8% ± 10.8%) (**Figure 1A)**. The overlap between infection- and isolate-responsive DEGs was extensive, with nearly all up-regulated (99.4%) and most down-regulated (93.3%) infection-responsive DEGs also significant for isolate-specific modulation in all hosts. This indicates that while each host has a common transcriptional program responding to *B. cinerea* infection, this response is substantially reshaped by pathogen genetic variation.

**Figure 1:**
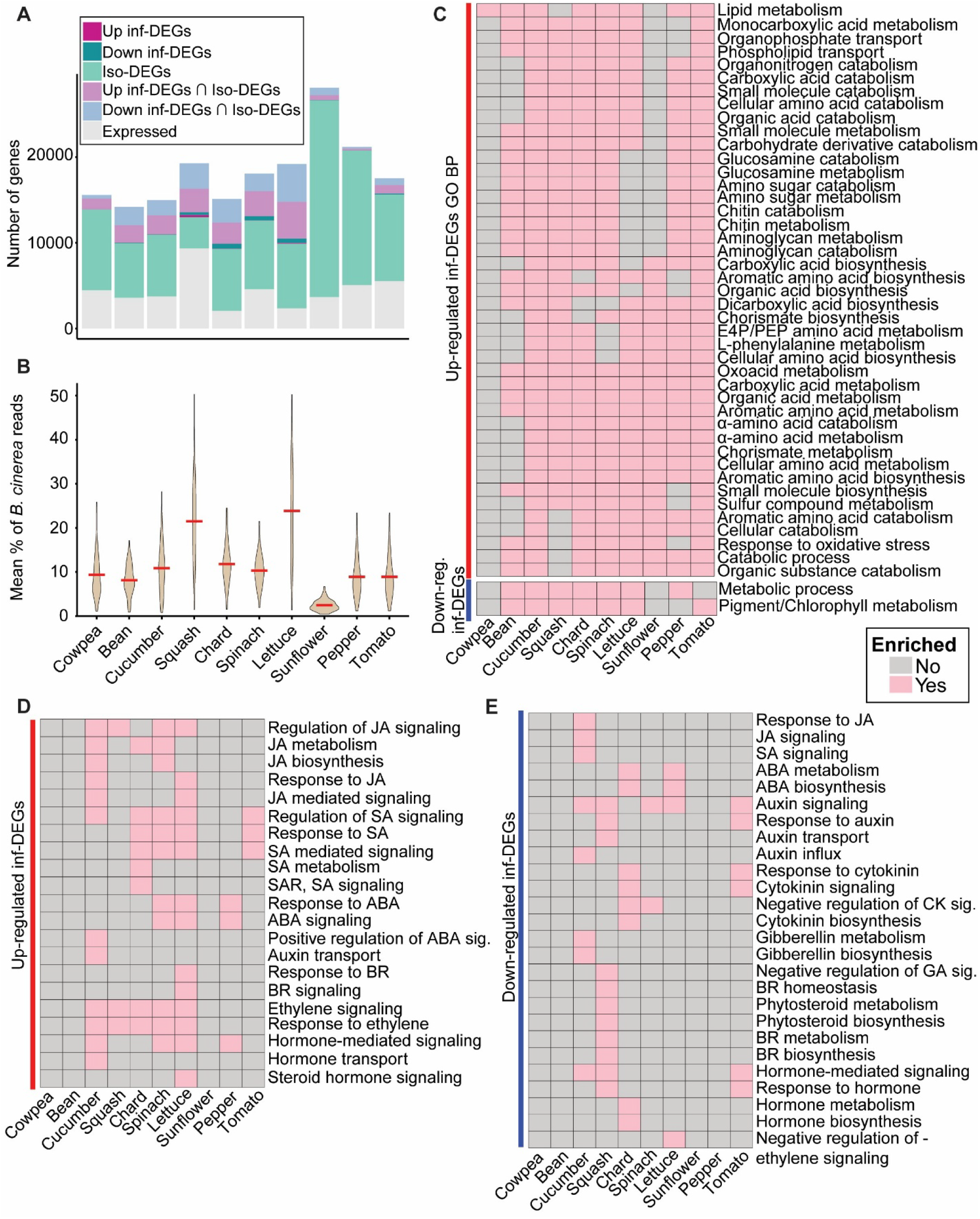
Host transcriptional responses and enriched functional pathways across ten eudicot species upon *B. cinerea* infection. **(A)** Summary of host gene expression responses. Bars indicate the number of total expressed genes, infection-responsive differentially expressed genes (inf-DEGs), and isolate-responsive DEGs (iso-DEGs). DEGs were identified using a generalized linear model (count ∼ infected + infected/isolate + tray + batch). Infection-responsive DEGs represent global response to infection (uninfected vs. infected; FDR ≤ 0.05, log₂FC > 1), irrespective of individual isolates. Isolate-responsive DEGs showed significant significant variation among isolates (Infection/Isolate; FDR ≤ 0.05). The overlap (∩) indicates genes significant for both categories. **(B)** Violin plots showing the mean percentage of *B. cinerea* reads, used as a proxy for fungal biomass. Values were calculated as the proportion of total RNA seq reads mapped to the *B. cinerea* genome relative to total reads mapped to both host and pathogen per sample. Red lines indicate medians. **(C)** Heatmap of Gene Ontology (GO) Biological Process terms significantly enriched (FDR ≤ 0.05) among up- and down-regulated infection-responsive DEGs shared across seven to ten host species. Pink = significant enrichment (FDR ≤ 0.05), gray = no significant enrichment. **(D-E)** Hormone-related GO enrichment among **(D)** upregulated and **(E)** downregulated infection-responsive DEGs. JA (Jasmonic acid), SA (Salicylic acid), ABA (Abscisic acid), BR (Brassinosteroid), SAR (Systemic Acquired Resistance), CK (Cytokinin), and GA (Gibberellin).

To understand what factors might influence the observed variability in the number of infection-responsive DEGs across hosts, we tested if the magnitude of the host transcriptional response correlates with pathogen biomass or virulence (measured as lesion size). Pathogen biomass i.e. percent of *B. cinerea* reads mapped was estimated as the proportion of total RNA-seq reads mapped to the *B. cinerea* genome relative to total reads mapped to the host and pathogen in that sample (Blanco-Ulate et al., 2014; Zhang et al., 2019; Singh et al., 2025). We first tested the relationship between fungal biomass and lesion size across hosts and found that the two were not significantly correlated **(Figure S2A)**. This indicates that visible disease severity is decoupled from the extent of early fungal colonization, potentially reflecting host-specific differences in defense or tolerance strategies. Given this decoupling, we tested each metric separately against the infection-responsive host DEGs. While fungal biomass and virulence both varied significantly among hosts (**Figure 1B; Figure S1**), virulence showed no significant correlation with the number of infection-responsive host DEGs (*r* = 0.08, *p* = 0.83) (**Figure S2B**). For example, lettuce and sunflower both developed large lesions; however, lettuce exhibited the strongest transcriptional activation, whereas sunflower showed one of the weakest responses. In contrast, fungal biomass was strongly correlated with the infection-responsive host DEGs (*r* = 0.82, *p* = 0.0038; **Figure S2C**). These results suggest that common host infection-response is associated with fungal biomass rather than lesion size. The disconnect between host transcriptional response and lesion indicates that host responses vary in effectiveness, allowing similar levels of fungal biomass to produce divergent disease outcomes. Notably, this connection only appears when investigating the common infection-response (infection-responsive DEGs) as the number of isolate-responsive host DEGs did not relate to the estimated biomass (**Figure S2E**).

### Shared metabolic remodeling across hosts but host-specific hormone responses

To identify defense responses to *B. cinerea* that appear conserved across eudicot hosts, we performed Gene Ontology (GO) enrichment analysis on up- and downregulated infection-responsive DEGs for each host and compared enriched processes among host species. Although no single GO term was significantly enriched across all ten hosts, 43 upregulated biological processes were shared among seven to nine hosts (**Figure 1C**). These broadly upregulated responses include primary metabolic processes such as catabolism of aromatic amino acid, organic acid, carbohydrate derivative, amino sugar, and lipid, with only limited enrichment of biosynthetic pathways such as chorismate, carboxylic/dicarboxylic acids, and aromatic amino acids. Additionally, cell wall-associated processes, including chitin and aminoglycan metabolism, were induced across seven hosts potentially reflecting a common activation of defense programs targeting fungal cell wall components. In contrast, analysis of downregulated pathways revealed a consistent suppression of chlorophyll/pigment metabolism. This agrees with the common observation that photosynthesis decreases soon after foliar infection by *B. cinerea* (BILGIN et al., 2010; Windram et al., 2012; Zhang et al., 2019; Soares et al., 2022; Cheaib and Killiny, 2025).

This shared pattern of metabolic reprogramming across multiple hosts upon *B. cinerea* infection raised a key question: is this shared defense framework orchestrated by a universal hormonal signaling network, or have different host plants evolved distinct regulatory strategies to achieve similar metabolic outcomes? Defense against necrotrophic pathogens is classically associated with JA and ET signaling (He et al., 2012; Yang et al., 2015; Hickman et al., 2019); with evidence also suggesting role of SA in local immunity against *B. cinerea* (Ferrari et al., 2003; Zhang et al., 2017). Specifically investigating these phytohormone pathways in our cross-species analysis revealed that hosts appear to contain species-specific signatures amongst these phytohormone signaling genes (**Figure 1D-E**). Spinach and lettuce exhibited the broadest hormonal reprogramming, inducing JA, SA, abscisic acid (ABA), and ET signaling genes. Cucumber and squash both induced JA and ET signaling genes, with cucumber additionally showing induction of ABA signaling genes. Chard and tomato predominantly induced SA signaling genes, whereas pepper only showed induction of ABA signaling genes. Conversely, hormone-related terms among downregulated genes largely reflected suppression of growth-associated pathways. Auxin signaling genes were repressed in cucumber, squash, spinach, lettuce, and tomato, while cytokinin signaling genes were suppressed in chard and tomato. Additional downregulation was observed for gibberellin (GA) signaling genes in cucumber and brassinosteroid (BR) genes in squash. Cowpea, bean, and sunflower did not show significant enrichment for any hormone-related processes, suggesting limited or transcriptionally subtle hormonal modulation in these hosts. Together, these results indicate that although majority of hosts reprogram hormonal signaling genes during infection, the specific combinations of induced and repressed pathways are highly lineage dependent, reflecting the potential for wide variation in host hormonal signaling underlying a shared defense framework.

To understand how these metabolic and hormonal responses converge at the transcriptional level, we analyzed the enrichment of TF and protein kinase families among infection-responsive DEGs within each host species. Unlike the GO analysis, this enrichment used combined up- and downregulated infection-responsive DEGs to capture overall regulatory involvement. Among 65 enriched TF families, WRKY and MYB TFs were consistently enriched across all ten hosts, with AP2/ERF, bHLH, bZIP, MYB-related, and NAC families enriched in nine hosts (**Figure S3A**). These results align with previous studies demonstrating roles of the ERF (Moffat et al., 2012), WRKY (Birkenbihl et al., 2012; Liu et al., 2015; Liu et al., 2017; Chen et al., 2021), MYB (Mengiste et al., 2003; Ramírez et al., 2011), bHLH (Ullah et al., 2022), and NAC (Wang et al., 2009; Windram et al., 2012) families in *B. cinerea* defense. Similarly, Receptor-like kinases (RLKs) and calmodulin-dependent protein kinase (CAMK) families were enriched in seven to nine species, pointing to their shared roles in initial pathogen perception and Ca²⁺-mediated signal transduction (**Figure S3B**). Interestingly, while many RLKs subclasses (e.g., *RLK-Pelle_LRR-XI-1*) were broadly enriched across most hosts, others were restricted to few species (*RLK-Pelle-LRR-XIII)* (**Figure S3B**). This observation suggests that while the overall signaling architecture involves similar gene families, specific components within those families appear to have diversified across evolutionary lineages. Together, these results suggest that plants have sets of conserved and lineage specific responses to *B. cinerea* that are controlled by a similarly changing set of regulatory machinery.

### Rapid evolution and functional divergence reshape conserved defense response across eudicots

To directly compare host transcriptional responses to *B. cinerea* infection across eudicots, we used OrthoFinder (Emms and Kelly, 2019) to group all protein encoding genes from the ten hosts into orthogroups, defined as sets of genes descended from a single ancestral locus. Direct comparison of the same orthogroups allowed us to better parse shared versus diverging aspects of the response. We identified 9815 core orthogroups present in all ten hosts, representing the shared ancestral gene set of these eudicots (**Figure S4A)**. Integrating these orthogroups with infection-responsive DEGs identified 147 orthogroups that responded to *B. cinerea* in at least eight host species, including a core set of 12 orthogroups that were infection responsive across all ten hosts (**Figure 2A and Figure S4B**). Among the 12 core infection-responsive orthogroups, five contained predominantly upregulated DEGs and encompassed potential defense/stress related genes such as *arogenate dehydratase 3* (phenylalanine biosynthesis; (Para et al., 2016)), *endochitinase* (cleavage of fungal chitin; (Cheng et al., 2017; Vaghela et al., 2022)), *WRKY75* (Chen et al., 2021), *cytochrome b561* (iron and ascorbate reductases; (Lu et al., 2014)), and *plastid movement impaired 1* (a coiled coil protein that alters plastid movement; (Didelon et al., 2020)) (**Table S3**). The remaining seven core infection-responsive orthogroups displayed mixed transcriptional directionality, containing both up and downregulated DEGs along with non-responsive genes. These orthogroups include genes annotated as *oligopeptide transporter 4*, *acidic endochitinase SP2*, *formin*, *CBL-interacting protein kinase 11* (*CIPK11*), *UDP-glycosyltransferase 71E1*, and two uncharacterized genes. Although the transcriptional direction of these orthologs differed among hosts, their consistent differential expression across eudicots suggests they potentially influence the *B. cinerea* response. This indicates that for some ancestral gene families, being a core/conserved responding orthogroup refers to a shared responsiveness to the pathogen rather than a uniform transcriptional direction, possibly reflecting functional diversification within a conserved defense framework.

**Figure 2:**
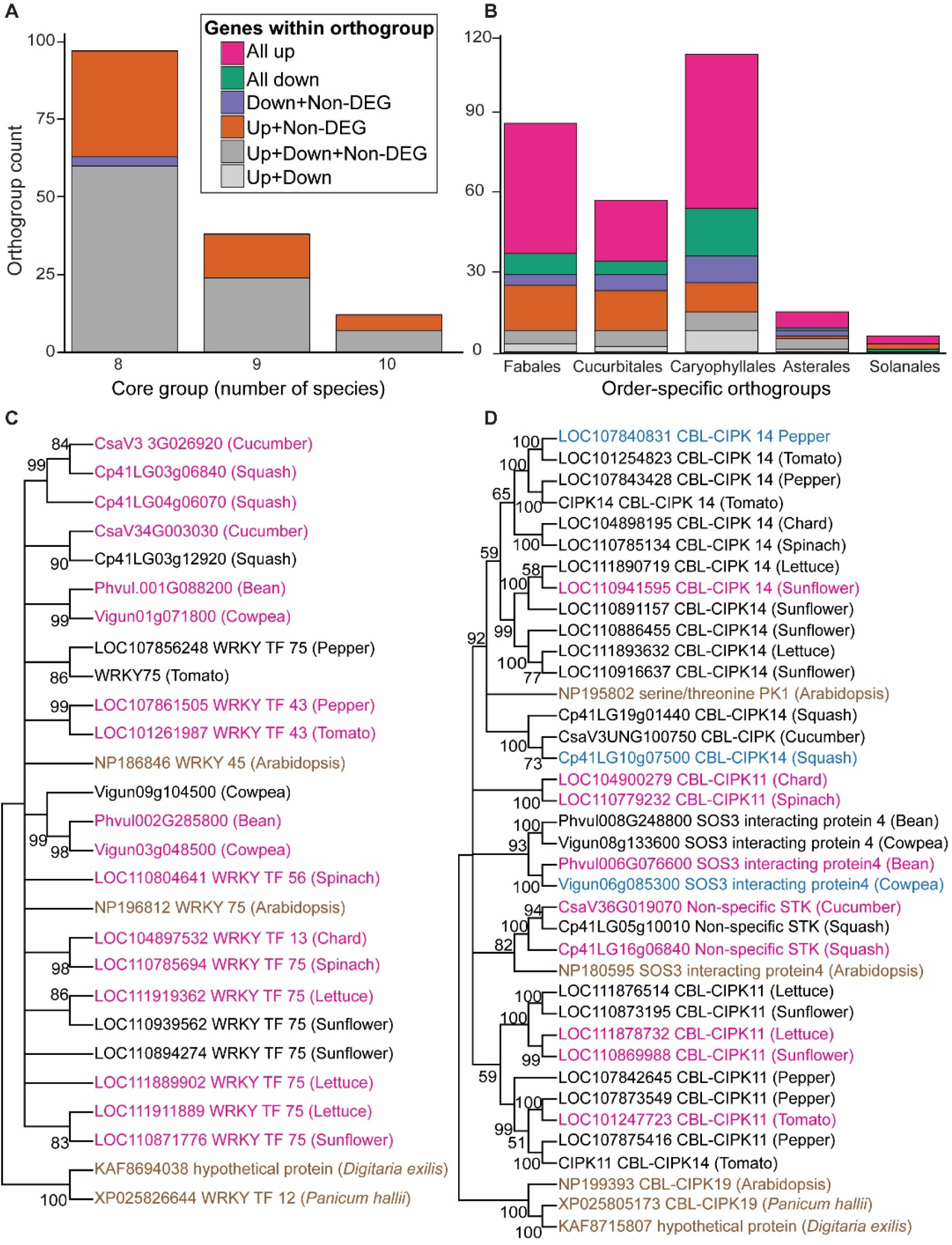
Core and order-specific infection-responsive orthogroups define conserved and divergent host responses. **(A-B)** Distribution and expression directionality of **(A)** core orthogroups shared across eight to ten host species and **(B**) order-specific orthogroups that are shared by both species within a given taxonomic order. Bar colours denote the directionality of gene expression responses within each orthogroup, as indicated in the key. **(C-D)** Neighbour-joining phylogenetic trees of two core orthogroups that are infection-responsive across all ten eudicot hosts: **(C)** *WRKY75* (OG0002688) and **(D)** CBL-interacting serine/threonine-protein kinase 11 (*CIPK11*, OG0001356). The trees were constructed in MEGA12 (Kumar et al., 2024)with 1,000 bootstrap replicates. Orthologs from *Arabidopsis thaliana* was included as functional anchors, and the trees were rooted using *Digitaria exilis* and *Panicum hallii* as monocot outgroups. Bootstrap support values are shown at each branch; branches with < 50% support collapsed into polytomies. Gene names are coloured according to their expression status: pink for upregulated, blue for downregulated, and black for non-differentially expressed genes. Reference genes from *A. thaliana* and monocot outgroups are shown in brown.

This regulatory divergence raises a key question: How have lineage specific gene duplications, losses, and regulatory changes reshaped transcriptional responses within these core orthogroups that are conserved across all hosts? To investigate this, we reconstructed phylogenies of two representative core infection-responsive orthogroups: the largely upregulated *WRKY75* and the directionally variable *CIPK11.* Phylogenies were built using all gene members, regardless of expression, within these orthogroups from each host plant species. Arabidopsis homologs were included as functional anchors and monocot sequences from *Digitaria exilis* and *Panicum hallii* as outgroups. In both orthogroups, gene trees deviated from the established species phylogeny, indicating numerous duplications and gene losses across the species within these orthogroups (**Figure 2C-D)**. This level of duplication and gene loss is often found in genes rapidly responding to pathogen or other stress. *WRKY75* orthogroup, which includes *A. thaliana AtWRKY75* known to be involved in the *B. cinerea* response (Chen et al., 2021), showed species-specific duplication and divergence in transcriptional regulation. While most members of this orthogroup were significantly upregulated upon infection, the expression of paralogs within a given species differed (**Figure 2C)**. For instance, pepper and tomato each possess two copies, but only one was infection responsive. Squash and cowpea each harbored three copies, with two being upregulated while in sunflower only one of three paralogs was infection responsive. The transcriptional response of *CIPK11* ortholog showed even more heterogeneity: pepper and cowpea orthologs were downregulated, squash contained two paralogs with opposite transcriptional responses (one up, one down), and other hosts retained at least one upregulated copy (**Figure 2D)**. These patterns indicate that even within core infection-responsive orthogroups, the individual genes are undergoing rapid evolution through lineage specific duplications and gene losses. This divergence in transcriptional regulation underscores the evolutionary plasticity of signaling networks in adapting to specific host-pathogen interactions.

### Order-Specific Orthogroups Reveal Specific Evolution in Immune Responses

To identify lineage-specific defense strategies beyond the core pathways shared across eudicots, we investigated order-specific orthogroups: gene families restricted to a single eudicot order. Whole-genome comparison identified 6,788 orthogroups confined to a single plant order, ranging from 881 in the Cucurbitales to 2,234 in the Fabales (**Figure S4A**). When integrated with infection-responsive DEGs, 276 orthogroups were responsive to *B. cinerea* infection in both species within their respective order, ranging from 6 in Solanales to 112 in Caryophyllales (**Figure 2B**). An additional 1,146 order-specific orthogroups were infection responsive in only one of the two species per order, indicating that infection-related transcriptional responses diverge even among closely related hosts.

Among these 276 shared order-specific infection-responsive orthogroups, approximately half of the orthogroups contained genes that were consistently upregulated in both hosts, whereas very few were consistently downregulated (**Figure 2B; Table S4**). These commonly induced order-specific orthogroups represent gene families uniquely deployed by each order during infection. A major caveat in interpreting these results is that orthogroup annotation is largely based on Arabidopsis homologs; thus, while gene names may appear similar to those in other orders, these orthogroups are lineage-restricted gene families. Therefore, annotations should be viewed as broader gene family annotations and not specific orthology identity. Caryophyllales specific upregulated orthogroups included receptors and signaling components, such as *wall-associated receptor kinase 11*, *RPM1-interacting protein*, *auxin-binding protein ABP19b* family, and *ERF115*, as well as enzymes for the biosynthesis of phenylpropanoids, including *chalcone synthase*, *caffeoyl-CoA O-methyltransferase*, and *flavanone 3-dioxygenase*. These suggest a possible defense module coupling early perception and hormone signaling with reinforcement of the cell wall. Cucurbitales specific orthogroups contain hormone signaling genes such as *TIFY10* family (JA signaling), *ERF5* (ET signaling), and *9-cis-epoxycarotenoid dioxygenase* (ABA biosynthesis). These signaling genes were paired with metabolic and structural proteins such as *extensin*, *glycosyltransferase*, *glyoxalase*, *VQ-domain*, *LEA2 domain*, *carbonic anhydrase*, and *isoflavone reductase* protein. Fabales order-specific orthogroups include *WRKY33*, *MYB14*, *MYB62*, and *heat shock factor* TFs, along with *dirigent-like* and *peroxidase superfamily* genes, and Ca²⁺-signaling components. Asterales specific orthogroups, exhibited induction of *costunolide synthase* enzymes associated with sesquiterpene lactone biosynthesis, known antifungal metabolites (Chadwick et al., 2013) along with the immune regulators *DMP3/DMP5* (basal immunity) and *PUB24* (negative regulator of pattern-triggered immunity; (Trujillo et al., 2008)). Solanales induced orthogroups were annotated as *Kunitz trypsin inhibitors* (do Amaral et al., 2022), *PHOS32/34* phosphorylation targets (Merkouropoulos et al., 2008), and *BURP-domain* proteins (Yu et al., 2022), indicating pathogen-induced protease inhibition and cell wall remodeling. These order-specific responses suggest that each lineage has a distinct set of defense components from signaling to output. Together, these order-specific responses show that, although eudicot hosts share a broadly conserved immune framework, each evolutionary lineage has independently diversified specific gene families to fine-tune its defense strategies.

Finally, at the most specialized layer, we examined species-specific orthogroups. These are defined as gene families that are genetically unique, with no detectable orthologs in any of the other nine tested hosts. Out of 7,402 such orthogroups identified across the ten genomes, 1,137 were responsive to infection in their respective host (**Figure S4B)**. GO enrichment analysis of these species-specific infection-responsive orthogroups did not reveal any significant enriched functional categories, both within and across species. This lack of enrichment within species suggests that these genes are not converging on a few common pathways but are instead drawn from diverse functional pools from whole-genome evolutionary events like duplications. Similarly, the absence of common GO terms across species suggests that each host has evolved a distinct repertoire of defense-related genes tailored to its own physiology and ecological niche. In total, this shows that the host infection-responsive DEGs identify a blend of genes that are shared across all hosts, order-specific, and species-specific.

### Diversified Host Plant Responses to Pathogen Variation

In the above sections, we focused solely on the host genes that had a common response to *B. cinerea* as a species, i.e. same across isolates (infection-responsive DEGs). However, this common response is layered upon a much larger landscape of isolate-responsive DEGs. In every host, most of the transcriptome was significantly modulated by the isolate-dependent response, showing that pathogen genetic variation plays an important role in shaping host transcriptional responses across all hosts (**Figure 1A**). We therefore asked whether the diverse plant hosts have a common response to the *B. cinerea* isolate diversity or if each host responded differently to the pathogen genetic variation. To provide a directly comparable framework across hosts, we focused on 803 single-copy orthologs defined as genes with a single homolog in all ten host species. These provide a common reference set across all the hosts to quantify the similarity or difference in responses to the pathogen genetic variation. Expression data for these genes were used to perform principal component analysis (PCA) of host transcriptional responses to the 72 *B. cinerea* isolates, including uninfected controls as reference.

Across all ten hosts, the first principal component (PC1) separated infected from uninfected samples (**Figure 3A**). This axis captured the primary transcriptional shift associated with infection in every species. Importantly, this pattern was conserved across hosts, indicating that infection induces a broadly similar direction of transcriptional change in eudicots. We treat this PC1 axis as a comparative response framework that summarizes shared host transcriptional reprogramming. To interpret the biological processes associated with this axis, we examined genes with the strongest positive and negative PC1 loadings (**Figure S7A**). Genes contributing positively to PC1 were related to stress and defense processes, including ABA regulator *AIP2*, programmed cell death-associated *CAD1*, and glycosylation enzymes (Morita-Yamamuro et al., 2005; Zhang et al., 2005; Chen and Cheng, 2024). Conversely, genes contributing negatively to PC1 were associated with growth, development, and ribosomal biogenesis such as dynamin-like *ARC5, RNA helicase 50,* and ribosomal factor *RsgA1* (Gao et al., 2003; Motohashi et al., 2007; SCHMITZLINNEWEBER and SMALL, 2008; Qiao et al., 2013; Rocchio et al., 2019; Ru et al., 2021; Fuentes-Terrón et al., 2024; Wang and Wu, 2025). This pattern is consistent with the shift away from growth associated processes toward stress-related transcriptional programs during infection (Karasov et al., 2017; Monson et al., 2022).

**Figure 3.**
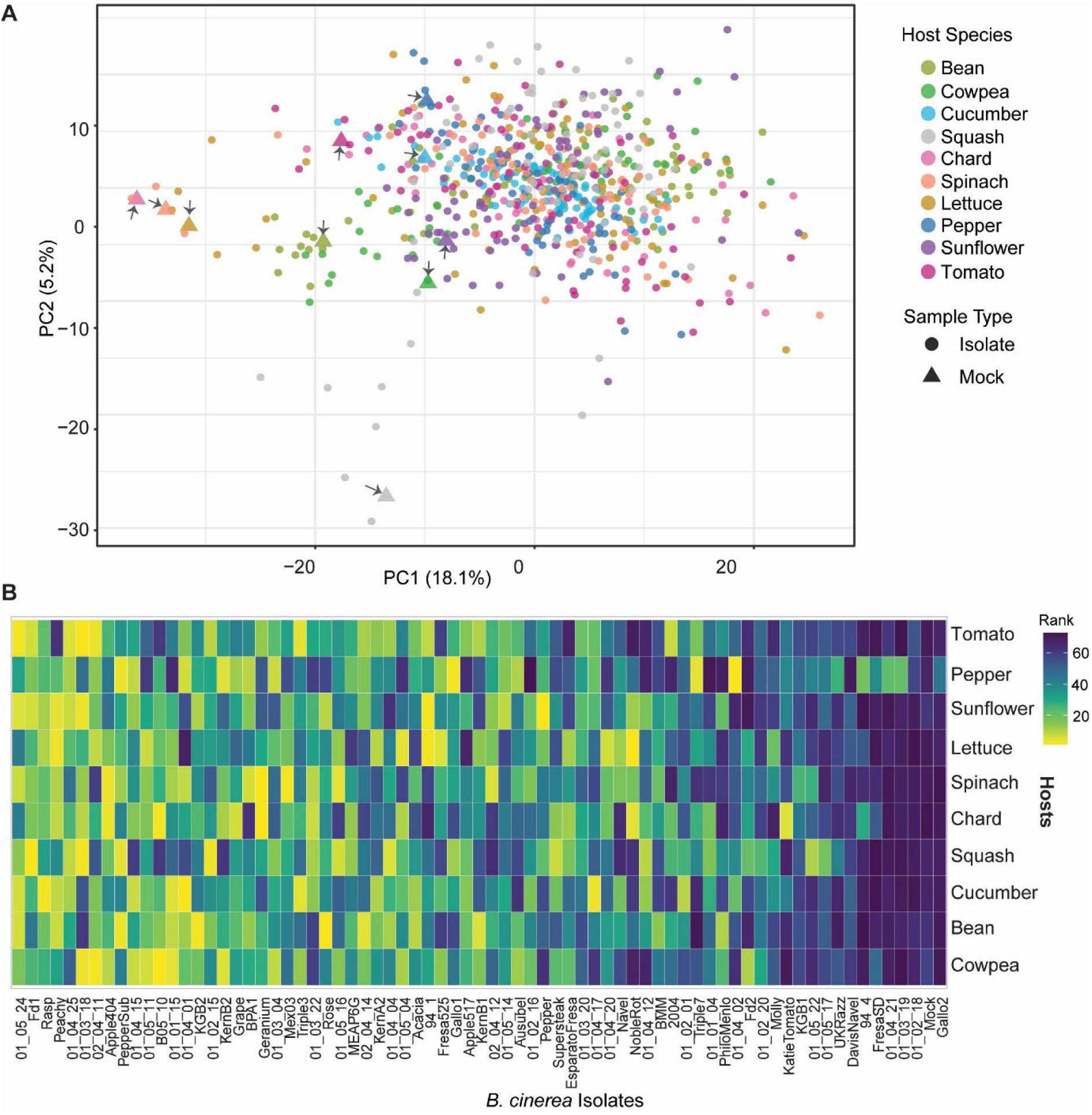
Isolate-driven transcriptional responses across hosts. (**A**) Principal component analysis (PCA) of host transcriptional responses based on the expression of 803 single-copy orthologs shared across all ten eudicot hosts. To normalize for interspecific scale differences, expression values were z-scaled within each host prior to analysis. Points are colored by host species and shaped by treatment (circles = *B. cinerea* infected; triangles = mock). Mock samples are additionally highlighted with arrows. The percentage of variance explained by each principal component is indicated on the axes. (**B**) Rank heatmap of *B. cinerea* isolates based on their Principal Component 1 (PC1) scores, which capture the primary axis of transcriptional variation. Isolates (columns) are ordered by their average rank across all hosts. Host species (rows) are arranged according to their phylogenetic relationships. The color scale represents the rank of each isolate within a host, where 1 (yellow) corresponds to the highest PC1 score (strongest transcriptional response) and higher ranks (purple) indicate weaker responses closer to mock controls.

While PC1 represents a common host response vector to the pathogen’s infection, it wasn’t clear if each host uses this common response in the same way across the diverse isolates. In other words, does the same pathogen isolate elicit a similar response across different hosts, or is the perception of pathogen variation unique to each host-pathogen combination? To test this, isolates were ranked within each host based on their PC1 score. If hosts responded similarly to pathogen genetic variation, the rank order of isolates would be consistent across species. Instead, isolate rankings shifted markedly among hosts: isolates that elicited strong transcriptional response in one species often induced moderate/weak response in another (**Figure 3B**). This lack of concordance indicates that the host’s transcriptional response to pathogen genetic variation is not intrinsic to the isolate but is defined by the specific host-isolate interaction. A small subset of isolates (Gallo2, 01_02_18, 01_03_19, 01_04_21, FresaSD, 94_4, DavisNavel, UKRazz, 01_05_17) maintained relatively stable rank order across hosts, with consistently weak transcriptional responses (**Figure 3B**). Lesion phenotyping showed that these isolates were not broadly avirulent **(Figure S8)**. To distinguish between low overall virulence and host-restricted pathogenicity, we compared each isolate’s mean PC1 score with the coefficient of variation (CV) of lesion sizes across hosts (a measure of host range). This showed that many of these weakly-responsive isolates tend to exhibit high host specificity, causing strong disease in limited number of hosts while remaining weak in others **(Figure S8**). In contrast, most of the isolates that had variable PC1 ranking across hosts were more likely to function as broad-host-range generalists (low CV) (**Figure S7B**).

Finally, we asked if the magnitude of this shared transcriptional response (PC1) predicts the disease outcome (lesion size) within individual hosts, and whether this relationship is similar across hosts. If the immune response functions similarly across eudicots, we expect a consistent correlation between host transcriptional response and disease severity. Instead, correlations between PC1 scores and lesion sizes revealed host-specific patterns. Moderate positive correlations were observed in the cucurbitales, caryophyllales, and asterales, whereas no significant association was found in the fabales and solanales (**Figure S5**). Interestingly, removing the subset of weakly-responsive isolates altered these correlations, indicating that a small number of isolates can disproportionately influence the relationship between host transcriptional activation and disease severity in a host dependent manner (**Figure S6**).

Together, these findings demonstrate that the magnitude and direction of host transcriptional responses are jointly shaped by the specific host-pathogen isolates combination. While eudicot hosts share a common direction of transcriptional reprogramming during *B. cinerea* infection, the deployment and phenotypic relevance of this response vary among hosts. This highlights the highly context-dependent nature of eudicot-*B. cinerea* interactions, where conserved defense programs are dynamically reprogrammed by pathogen genetic variation.

### Host Transcriptional Responses to *B. cinerea* Toxins Are Species-Specific and Not Evolutionarily Conserved

To defend against the necrotrophic attack of *B. cinerea*, host plants must cope with the pathogen’s functionally overlapping virulence mechanisms, including the key phytotoxins *Botrydial* (*BOT*) and *Botcinic acid* (*BOA*) (Leisen et al., 2022), which contribute to tissue damage and disease progression. While these toxins are well established components of the *B. cinerea* infection process, it remains unclear whether hosts have a conserved transcriptional response to these toxins or whether these responses are lineage specific. Addressing this question from the host perspective is critical for understanding how phylogenetically diverse hosts respond to these common virulence factors.

To examine how hosts transcriptionally respond to toxin exposure, we leveraged natural variation in *BOT* and *BOA* pathway expression across 72 *B. cinerea* isolates (Singh et al., 2025). Previous analyses showed that genetic variation in these virulence mechanisms is largely independent across isolates, providing a controlled source of differential toxin challenge (Deighton et al., 2001; Siewers et al., 2005; Singh et al., 2025). In many plant-pathogen systems, host genes involved in toxin perception, detoxification, or downstream signaling are transcriptionally responsive to toxin exposure (Rossi et al., 2011; Walter and Doohan, 2011; Lorang et al., 2012). We therefore hypothesized that host genes whose expression correlates with fungal toxin gene expression could represent potential toxin responsive host genes. By comparing these responses across ten phylogenetically diverse eudicot species, we tested whether host transcriptional responses to *BOT* and *BOA* reflect conserved defense mechanisms or lineage-specific strategies.

We first assessed whether the host’s capacity to mount a transcriptional response track with the fungal toxin expression across species. Although fungal *BOT* and *BOA* biosynthetic genes were expressed during infection in all hosts, albeit with quantitative variation (**Figure S9A**), host transcriptional responses varied markedly. High levels of fungal toxin expression did not necessarily trigger a robust host transcriptional response (**Figure 4B and S9B**). For example, lettuce and sunflower both experienced high fungal toxin expression, yet only lettuce showed a significant set of host genes correlated with toxin expression, whereas sunflower showed little to no transcriptional association. This implies that sunflower may lack the specific molecular components required for toxin perception or downstream signaling, potentially contributing to its high susceptibility (**Figure S1**). Across the ten hosts, toxin-associated transcriptional responses were unevenly distributed. Four species (cowpea, bean, lettuce, pepper) showed significant host gene correlations with both *BOT* and *BOA* expression, three species responded only to *BOT* (cucumber, squash, and chard), and the remaining species showed minimal or no detectable transcriptional association with either toxin (**Figure S9B**). These patterns indicate that the capacity to detect or transcriptionally respond to *B. cinerea* toxins is not a conserved feature of eudicot immunity but instead reflects lineage-specific differences.

**Figure 4:**
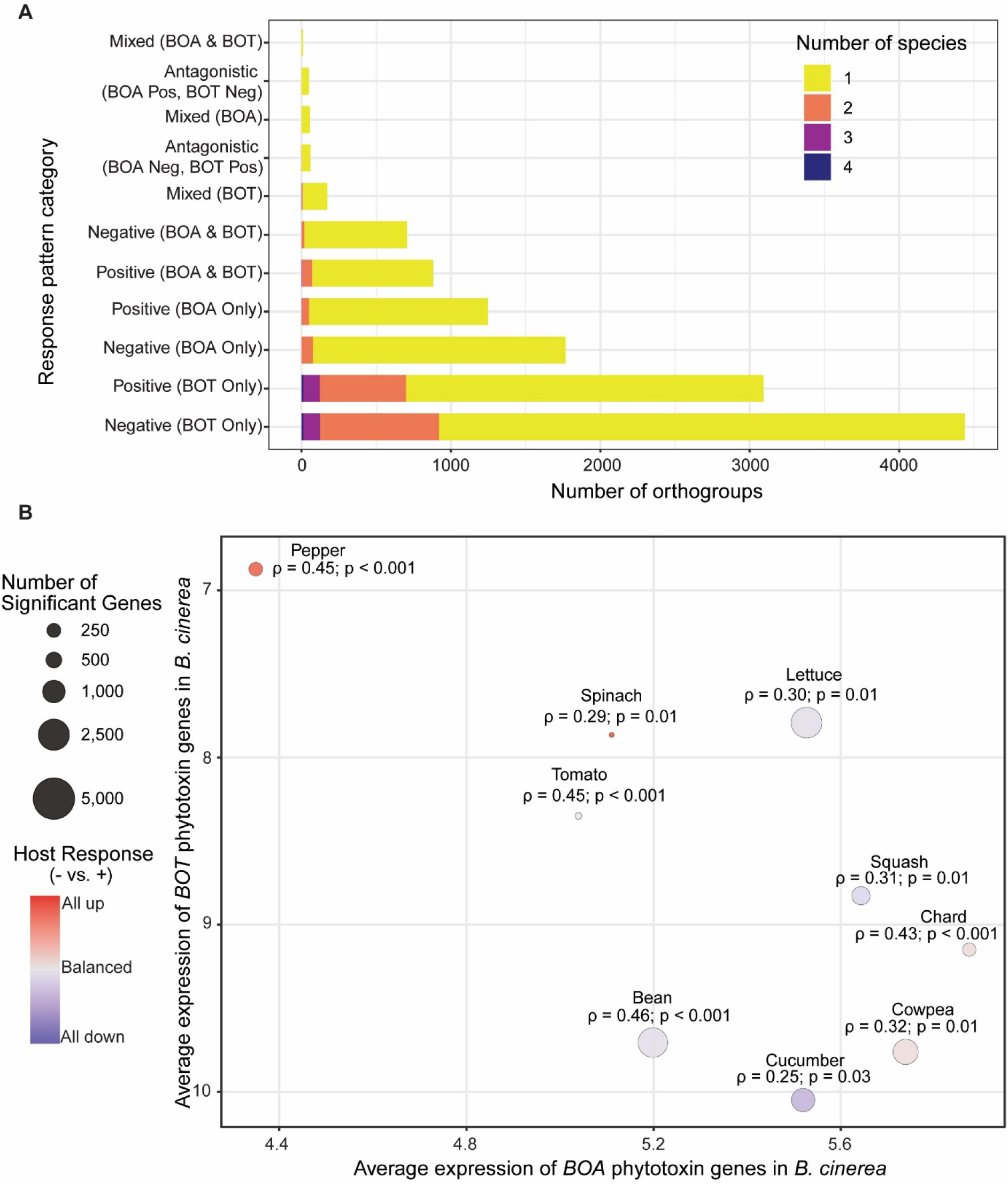
Host-specific transcriptional associations between host genes and pathogen phytotoxins. **(A)** Distribution of significant correlation patterns between host gene expression and pathogen phytotoxin biosynthetic gene expression for *Botrydial* (*BOT*) and *Botcinic acid* (*BOA*), summarized at the orthogroup level. Bar length indicates the total number of orthogroups exhibiting a given correlation pattern, while stacked colors denote the number of host species in which that pattern was observed. The mixed categories represent orthogroups exhibiting inconsistent directionality, specifically containing member genes with both positive and negative correlations for *BOA* and/or *BOT*. **(B)** Relationship between pathogen phytotoxin expression and host transcriptional responsiveness in each species. Each point represents a host species, positioned according to the average induced expression of *BOA* (x-axis) and *BOT* (y-axis) biosynthetic genes in *B. cinerea*. Average toxin expression was calculated across all genes within each biosynthetic cluster, *BOA* (*Bcin01g00010*-*Bcin01g00160*) and *BOT* (*Bcin12g06370-Bcin12g06410*), using isolate-level expression values. The size of each point is proportional to the magnitude of the host transcriptional response (number of significantly correlated genes), and point color reflects the directionality of the response, ranging from positive (red) to negative (blue) correlations. Spearman’s rank correlation coefficient (ρ) and associated *p*-value for the relationship between *BOA* and *BOT* induction are shown for each host. Sunflower is not shown due to absence of significant toxin-associated host genes. Individual host-level correlation plots and toxin expression data are provided in **Figure S9.**

To determine whether any toxin-responsive host programs were conserved across host species, we integrated significantly toxin correlated host genes with orthogroup annotations. The majority of toxin-correlated orthogroups were restricted to one or two species, indicating limited conservation (**Figure 4A**). The largest overlap was observed for BOT-associated responses, where 11 orthogroups were shared across four hosts, including *CML41*, *S-adenosylmethionine synthase 2*, *phospho-2-dehydro-3-deoxyheptonate aldolase 2*, *6-phosphogluconate dehydrogenase*, *coatomer subunit α-1*, *septum-promoting GTP-binding protein 1*, *TIM23-2*, *arabinogalactan O-methyltransferase 1-like*, and *cytochrome P450*. BOA-associated responses were far less conserved, with only a single orthogroup (*GH3.1*, indole-3-acetic acid amido synthetase) shared among three hosts. Three orthogroups (*cationic peroxidase 1-like*, *dihydroflavonol 4-reductase*, and *TIFY10b*) were correlated with both toxins across three species (**Table S5**). Thus, even among hosts that exhibited transcriptional responses to toxin exposure, the underlying gene families were largely distinct.

Together, these results demonstrate that host transcriptional responses to *B. cinerea* toxins are highly species-specific rather than being broadly shared. While fungal *BOT* and *BOA* biosynthetic pathways are broadly expressed across hosts, plant responses to these toxins vary substantially in magnitude and gene identity. This lack of shared, toxin-specific host responses suggests two possibilities: 1) the molecular targets of *BOT* and *BOA* are different in each host, reflecting divergent evolutionary trajectories, or 2) the actual targets are not transcriptionally regulated by the toxins in all hosts and the identified correlated genes largely track host responses to toxin-induced stress rather than molecular targets of *BOT* or *BOA*.

## Discussion

Advancements in molecular and genomic tools have greatly expanded our understanding of plant-pathogen interactions, yet translating this knowledge into durable disease resistance remains challenging, particularly against pathogens with high genetic diversity and polygenic virulence strategies (Corwin et al., 2016). A key limitation is that most resistance mechanisms are characterized in a small number of model systems and then implicitly assumed to be broadly conserved. This assumption remains largely untested across evolutionary scales, especially for generalist pathogens that infect many host lineages. By using a large-scale co-transcriptomic approach across ten eudicot species and 72 *B. cinerea* isolates, we demonstrate that plant defense is not a single conserved pathway, but a mosaic of deeply conserved physiological outcomes executed through extensive, lineage-specific regulatory rewiring. Pathogen genetic diversity further modulates both the magnitude and direction of these responses, demonstrating that disease outcomes arise from specific host-by-isolate interactions rather than species-level averages.

### A Conserved Core with Lineage-Specific Solutions

The central goal of this study was to determine how conserved and divergent immune components are deployed when evolutionarily distant hosts are challenged by the same, genetically variable pathogen species. GO analyses identified that broad physiological outcomes of infection such as metabolic reprogramming, cell wall modification, and suppression of growth-associated processes are shared across eudicots. However, the genetic basis underlying these responses is highly divergent. Among nearly ten thousand orthogroups conserved across all ten hosts, only 12 were consistently responsive to *B. cinerea* across all hosts. This small core represents a limited set of defenses and signaling components, indicating that while the physiological strategy of responding to necrotrophic infection is conserved, the specific genetic regulatory toolkit underlying this is divergent. Interestingly, even these 12-core infection-responsive orthogroups showed evidence of rapid evolutionary turnover, characterized by lineage-specific gene duplications, losses, and transcriptional divergence among paralogs. This pattern suggests that even the conserved defense frameworks are not static but are continually reshaped through microevolutionary processes. Such rapid diversification is a hallmark of host-pathogen arms races, where recurrent selective pressure drives the diversification of defense-related gene families (Lažetić and Troemel, 2021). This diversification is happening so quickly that it would require much deeper trees, involving more species with measured pathogen responses, to track the ancestral state of the pathogen response and the evolutionary progression within these gene families.

In contrast to this limited conserved core, the majority of the host response was defined by order- and species-specific orthogroups. These lineage-restricted genes represent independently evolved defense modules, likely shaped by ecological and pathogen-driven selection unique to each lineage. Rather than relying on a universal regulatory architecture, hosts deployed distinct regulatory solutions to achieve similar defensive outcomes, for example through species-specific deployment of hormonal signaling pathways. Furthermore, while lineage-specific genes are often assumed to primarily encode specialized metabolism, we found that many order- and species-specific orthogroups identified here encode signaling and structural components, demonstrating that evolutionary innovation in plant defense extends beyond secondary metabolism. This lineage-specific strategy extends even to host responses against common virulence factors as host transcriptional responses to ubiquitous phytotoxins were largely lineage-specific rather than shared. Notably, even closely related species within the same order frequently exhibited divergent transcriptional responses, highlighting that defense diversification operates across multiple phylogenetic scales. Thus, the plant immune system appears to be built on a foundation of independent regulatory solutions converging on shared defense outcomes.

### The indispensable role of pathogen diversity in understanding host response

Our study highlights the importance of incorporating natural pathogen diversity when dissecting host-pathogen interactions. By using 72 genetically diverse *B. cinerea* isolates, we moved beyond traditional single-isolate studies and were able to separate host responses that are consistent across all pathogen isolates from those that depend on isolate-specific variation. Nearly three-quarters of expressed host genes showed significant isolate-dependent effects, indicating that host transcriptional responses are highly plastic and strongly shaped by pathogen genetic diversity. From the host’s perspective, *B. cinerea* therefore represents a genetically heterogeneous population rather than a uniform threat. This isolate-dependent variation has important implications for how plant immunity operates in natural environments, where host immune response reflects the integration of signals generated by multiple pathogen variants. Even within the shared axes capturing global infection host responses, we found that both the magnitude and direction of host transcriptional response varied significantly across host-by-isolate combinations. Distinct host species mount divergent responses to the exact same isolate; activating a robust defense in one instance while responding weakly in another, emphasizing that transcriptional outcomes are defined by specific host-isolate interactions rather than uniform species-level responses.

The biological basis of this extensive plasticity likely reflects the high sensitivity of the host immune system to the variable virulence repertoires deployed by *B. cinerea*, including quantitative differences in phytotoxins, detoxification enzymes, and cell wall-degrading factors (Colmenares et al., 2002; Pedras et al., 2005; Rowe and Kliebenstein, 2007; Pedras et al., 2008; Pedras et al., 2011; Schumacher, 2012; Viaud et al., 2016; Zhang et al., 2019; Singh et al., 2025). Rather than a fixed program uniformly triggered by infection, host immunity seems to be capable of detecting the specific molecular arsenal deployed by a given isolate and adjusting its response magnitude to match the immediate threat. This implies that the host possesses a much higher resolution for sensing pathogen variation than previously appreciated, effectively discriminating between distinct isolate-derived molecular patterns. Consequently, the generalist host response is a collection of highly specific responses tailored to the immediate molecular context of the infection. Fully elucidating this two-way communication at the molecular level specifically how hosts perceive and integrate these complex signals to fine-tune their transcriptional output and how fungal isolates perceive distinct host environments remains a critical next step.

### Conclusion and Future Perspectives

These findings provide a systems-level understanding of how plant immune systems are tuned across evolutionary scales, shedding light on why resistance mechanisms often fail to translate across species and why durable resistance against generalist pathogens remains difficult to achieve. Durable resistance will likely require multilayered strategies that integrate conserved basal defenses with mechanistically distinct, lineage-specific modules that are resilient to pathogen adaptation. Our results further raise key evolutionary questions: Do lineage-specific defense networks evolve in parallel with conserved immune mechanisms, or do they emerge through independent regulatory rewiring? What ecological or genomic factors drive these shifts across plant orders? Addressing these questions will require expanded co-transcriptomic analyses across additional species and deeper phylogenetic sampling, including early diverging land plants. Integrating spatially resolved transcriptomics with functional validation will be essential to connect transcriptional divergence with cellular mechanisms and to clarify how conserved and lineage-specific defenses are deployed at the host-pathogen interface.

## Materials and Methods

This study utilizes the same dataset previously described in (Singh et al., 2025), which investigated *Botrytis cinerea* infection strategies across multiple eudicot species. Relevant portions of the materials and methods are repeated here, with minor modifications, to ensure clarity and accessibility for readers.

### Plant Material and Growth Conditions

A phylogenetically diverse panel of ten eudicot species was selected for this study. This panel represent three clades (Asterids, Super-Asterids, and Rosids) across five orders, with two species chosen per order: Asterales (*Helianthus annuus* , sunflower; *Lactuca sativa*, lettuce), Solanales (*Capsicum annuum*, pepper; *Solanum lycopersicum*, tomato), Caryophyllales (*Spinacia oleracea*, spinach; *Beta vulgaris*, chard), Fabales (*Vigna unguiculata*, cowpea; *Phaseolus vulgaris*, bean), and Cucurbitales (*Cucumis sativus*, cucumber; *Cucurbita pepo*, squash). Within each order, the species sampled belonged to the same family. All selected species are annual diploids with available reference genomes and reported susceptibility to *B. cinerea*. Seeds were obtained from the USDA-GRIN, the UC Davis Tomato Genetics Resource Centre, the CGN (Netherlands), and the Michelmore lab (UC Davis). Plants were grown in controlled environment chambers maintained at 20°C under a 16-hour light and 8-hour dark photoperiod, with light intensity of 100-120 µmol m⁻² s⁻¹. This subset was chosen because these species generated reliable fungal reads at 48 hours post inoculation (hpi), were diploid, easy to grow under controlled conditions, and supported consistent *B. cinerea* infection necessary for large-scale co-transcriptome analysis. To ensure experimental feasibility and enable cross-species comparisons within the broader framework of the study, a single representative genotype per species was used (**Table S6**). The selected genotype for each species represented intermediate lesion development based on prior lesion phenotyping data (Singh et al., 2025).

### *Botrytis cinerea* Isolates and Inoculum Preparation

A collection of 72 genetically diverse *B. cinerea* isolates (**Table S7**), originally collected from 14 different plant species across multiple geographic locations was used in this study. This collection has been previously characterized on several eudicot hosts (Zhang et al., 2017; Soltis et al., 2019; Zhang et al., 2019; Caseys et al., 2021; Singh et al., 2025) and captures broad variation in virulence without showing significant population structure by either host species origin or geography (Soltis et al., 2019; Caseys et al., 2021).

For inoculum preparation, isolates were revived from glycerol stocks onto potato dextrose agar (PDA) and incubated at 21°C for two weeks. Conidia were harvested from full-grown plates, counted using a hemacytometer, and diluted in 50% grape juice to a final concentration of 10 spores/μL (Singh et al., 2025). Grape juice, containing a natural mix of sugars and micronutrients, was used as the inoculation medium to ensure uniform germination across isolates and reduce variation due to genetic differences affecting germination on single-sugar sources (Blakeman, 1975; Benito et al., 1998; Denby et al., 2004).

### Detached Leaf Assay and Sample Collection

Infections were performed using a detached leaf assay, which recapitulates whole-plant infection responses (Govrin and Levine, 2000; Mengiste et al., 2003; Denby et al., 2004; Sharma et al., 2005; Windram et al., 2012; Soltis et al., 2020). Fully expanded leaves from each host species were inoculated with 4 µL droplets of a *B. cinerea* conidial suspension (10 spores/µL) and control plants were inoculated with grape juice only. Inoculated trays were placed under constant light and covered with humidity domes. Each isolate by host genotype combination was replicated three times in a randomized complete block design. Tissue was harvested at 48 hpi by excising 1.2 cm diameter leaf disks centered on the inoculation sites, followed by immediate flash-frozen in liquid nitrogen. This time point was selected based on a pilot RNA seq experiment conducted at 24, 30, 36, and 48 hpi, which showed that 48 hpi captured both an early stage of infection and yielded the highest proportion of fungal mapped reads across all host species, ensuring sufficient *B. cinerea* transcript abundance for cross species comparisons (Singh et al., 2025).

### RNA-Seq Library Construction, Sequencing, and Mapping

A total of 2,190 RNA-seq libraries were prepared and paired-end sequenced on the Element Biosciences AVITI150 platform at UC Davis DNA Technologies Core (Singh et al., 2025). RNA-Seq libraries were prepared following previous protocols (Kumar et al., 2012; Zhang et al., 2017; Singh et al., 2025). In brief, mRNA was extracted from frozen tissue using the Dynabeads mRNA Direct purification kit (Invitrogen). First and second strand cDNA synthesis was performed using Superscript III (Invitrogen). Libraries were then fragmented, end-repaired, A-tailed, barcoded, and size-selected (∼300 bp) as described previously (Kumar et al., 2012; Zhang et al., 2017; Singh et al., 2025). Libraries were pooled, multiplexed in sequencing batches, and sequenced.

Raw reads (FASTQ) were demultiplexed by adapter index. Read quality was assessed using MultiQC v1.15 (Ewels et al., 2016), and adapters/low-quality bases were trimmed with Trimmomatic v0.39 (Bolger et al., 2014). Reads were first aligned to host reference genomes (**Table S1**) using HISAT2 v2.2.1 with modified parameters to allow mismatches at read ends (Kim et al., 2019). Unmapped reads were subsequently aligned to the *B. cinerea B05.10* reference genome (Van Kan et al., 2017). Gene counts were generated using SAMtools (Li et al., 2009) and summarized across transcript isoforms using custom R scripts to avoid redundancy.

### Transcriptomic Data Analysis

All analyses were performed in the R v4.5.0 environment. For *B. cinerea*, read filtering followed previously described pipelines (Singh et al., 2025). For host expression analyses, expressed genes were defined as those with counts per million (CPM) > 1 in at least 40 samples per host species, using the cpm function in edgeR package (Robinson et al., 2010). Counts passing this threshold were normalized by the Trimmed Mean of M-values (TMM) method. For each host species, gene-level expression was modelled using a generalized linear model (GLM) with a negative binomial link function from the glmmTMB package (Venables and Ripley, 2002):

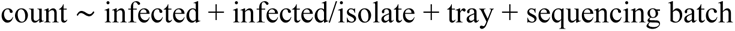

In this model, infected contrasts uninfected versus infected samples. The infected/isolate term represents isolate effects nested within infection, ensuring that isolate specific differences were modeled only among infected samples. Tray and sequencing batch accounted for experimental blocking and sequencing batch effects. Model-corrected means and standard errors for each transcript within each isolate-host combination were estimated using the emmeans package (Lenth, 2017). Wald χ² tests for model terms were obtained with the Anova function in car package, and *p* values were adjusted for multiple testing using the Benjamini-Hochberg procedure (Benjamini et al., 2001). Log fold changes for the infected main effect were estimated on the natural log scale and converted to log 2 scale.

Genes were designated as infection-responsive DEGs when the infected main effect was significant (FDR ≤ 0.05) with an absolute fold change greater than 1. These infection-responsive DEGs represent genes consistently reprogrammed across isolates and capture the average host transcriptional response to *B. cinerea* infection. In contrast, infected/isolate term identified genes whose transcriptional responses varied among isolates, thereby capturing isolate-dependent heterogeneity (called as isolate-responsive DEGs). These isolate-responsive DEGs were filtered solely based on FDR ≤ 0.05.

### Gene Ontology (GO), Transcription Factor (TF), and Protein Kinase (PK) enrichment analysis for infection-responsive DEGs

To identify overrepresented functional categories among infection responsive DEGs, GO enrichment analysis was performed separately for up- and down-regulated infection responsive DEGs for each host species independently. Analyses were conducted with the topGO package using species-specific GO annotations from corresponding GAF files. Enrichment was assessed for all three GO domains i.e. Biological Process (BP), Molecular Function (MF), and Cellular Component (CC) using the classic algorithm with Fisher’s exact test. Resulting p-values were adjusted for multiple testing within each species using the Benjamini-Hochberg procedure, with significance defined as FDR ≤ 0.05. To minimize redundancy, results were summarized using REVIGO (Supek et al., 2011). For cross-species comparison, enriched GO terms were converted into a binary presence-absence matrix and visualized as a heatmap using the pheatmap package in R (Kolde, 2010).

For regulatory gene families, combined infection responsive DEG sets (up- and down-regulated) were submitted to iTAK for TF and PK annotation (Zheng et al., 2016). The total number of predicted TFs and PKs in each host genome was retrieved from the same database to serve as the genomic background. Statistical enrichment of TF and PK families among infection responsive DEGs was then assessed for each species using two-tailed Fisher’s exact tests. Resulting *p*-values were adjusted using the Benjamini-Hochberg method, and families were considered significantly enriched at an FDR ≤ 0.05.

### Orthologous Gene Groups Analyses

Orthologous gene groups were identified by clustering the predicted proteomes of the ten host species using OrthoFinder v2.5.4 (Emms and Kelly, 2019) with default parameters. Based on phylogenetic distribution, orthogroups were classified into five categories: (i) core orthogroups, present in all ten hosts; (ii) clade-specific, restricted to either Asterids or Rosids; (iii) order-specific, restricted to a single order (Asterales, Caryophyllales, Solanales, Cucurbitales, or Fabales); (iv) species-specific, unique to one host; and (v) non-phylogenetically distributed, spanning multiple hosts without a clear phylogenetic pattern **(Figure S2A)**.

Infection responsive DEGs were then merged with these orthogroup classifications, and an orthogroup was considered significant in a given host if at least one gene was differentially expressed. Orthogroups were further categorized by transcriptional directionality, defined as uniformly upregulated (all_up), uniformly downregulated (all_down), or combination of up and down with non-DEGs. Functional annotation of orthogroups was performed by mapping member genes back to species-level GO and Pfam annotations.

### Correlation of Host Gene Expression with Pathogen Toxin Expression

To dissect host transcriptional responses specifically associated with *B. cinerea* phytotoxins, we performed a partial Spearman correlation analysis. A key challenge in this analysis is that toxin production is often correlated with overall virulence; thus, a host gene might appear correlated with toxin expression simply because it responds to the general intensity of infection. To account for this, we utilized general lesion-associated genes i.e., pathogen transcripts whose expression tracks overall virulence across isolates, as defined in previous study (Singh et al., 2025). By using the average expression of these genes as a covariate, we statistically controlled for the baseline effect of pathogen virulence, allowing us to isolate host responses that are specifically modulated by toxin pathway activity.

For each unique host-isolate combination, we correlated model-corrected host gene expression values (log₂ emmeans) with the average log₂ expression of genes within the *Botrytidal* (*BOT; Bcin12g06370*–*Bcin12g06410*) and *Botcinic acid* (*BOA; Bcin01g00010*–*Bcin01g00160*) biosynthetic cluster. The partial correlation was computed using the pcor.test function in the *ppcor* R package, with the general lesion-associated gene expression included as the controlling variable. For each host gene, we calculated Spearman’s rank partial correlation coefficients (ρ) and p-values for both *BOA* and *BOT*. P-values were adjusted for multiple testing separately within each host using the Benjamini-Hochberg (BH) method. Genes with FDR ≤ 0.05 were considered significant, with the sign of ρ indicating positive or negative association with toxin expression.

To identify evolutionarily conserved responses, all significantly correlated host genes were mapped to their respective orthogroups. Orthogroup-level responses were categorized based on the consistency of transcriptional directionality across species. Orthogroups were classified based on transcriptional directionality across member genes and species i.e. BOT-specific, BOA-specific, or dual-responsive (significant for both). Dual-responsive genes were further characterized as concordant (same direction of correlation for both toxins) or antagonistic (opposite directions). Orthogroups were designated as mixed if they exhibited inconsistent directionality, specifically containing genes with both positive and negative correlations for *BOA*, and/or both positive and negative correlations for *BOT*.

## Data, Materials, and Software Availability

All data supporting the findings of this study are included in the main article and/or the Supplementary Information. The RNA-seq data have been deposited in the NCBI Sequence Read Archive (SRA) under BioProject ID [PRJNA1217477] and will be made publicly available upon publication.

## Acknowledgements

This work was supported by the NSF award IOS 2020754 to DJK. We thank all members of the Kliebenstein lab for their invaluable assistance and help throughout the course of this large-scale experiment. We thank Michelmore lab (UC Davis) for providing lettuce seeds.

## Authors contributions

D.J.K.: designed research; R.S., A.J.M., C.T., and C.C.: performed research; R.S.: analyzed data; and R.S.: Writing - original draft, D.J.K.: Writing - review and editing.

---

**Figure S1.**
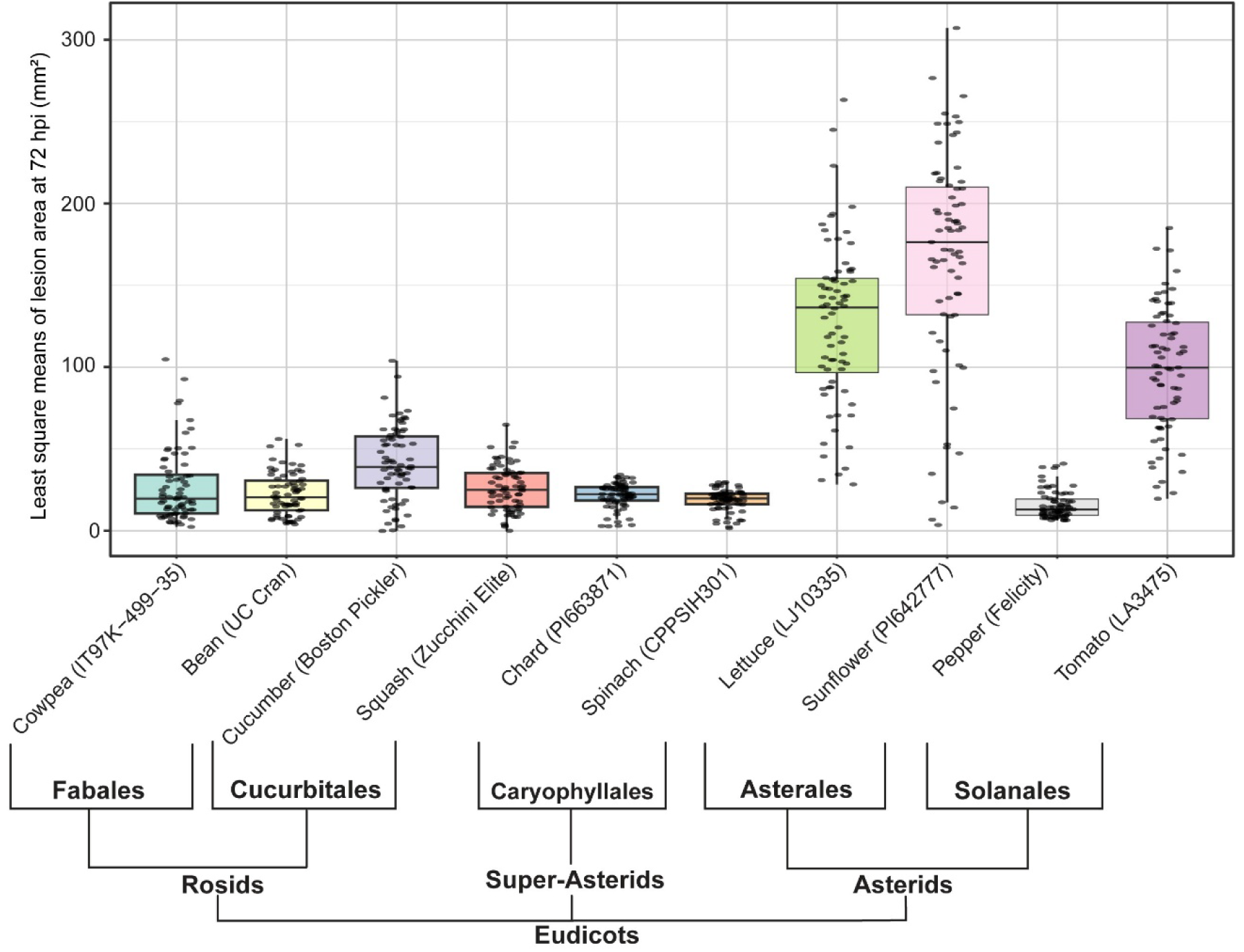
Average lesion size across ten selected eudicot hosts at 72 hours post inoculation. Boxplots show least square means of lesion area (mm²) caused by *Botrytis cinerea* across 10 host genotypes (shown in brackets) representing diverse eudicot species. Each point represents an individual isolate. Boxes indicate the interquartile range (IQR) with the median line, and whiskers extend to 1.5 x IQR. The phylogenetic relationships of the host species are indicated below, grouped by order and higher clades within the eudicots.

**Figure S2:**
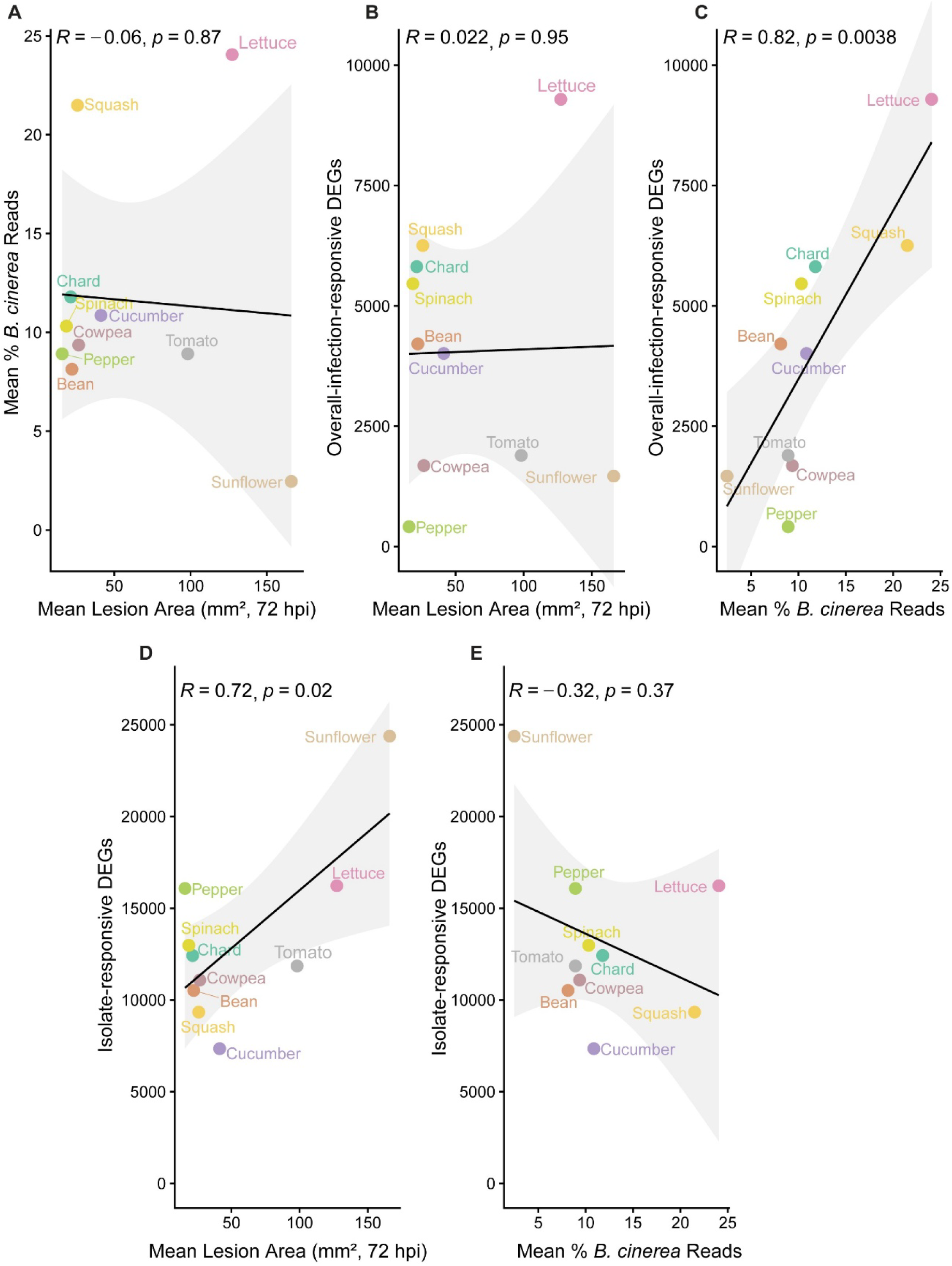
Correlation between fungal biomass, lesion size, and the number of differentially expressed host genes (DEGs) across ten eudicot species infected with *Botrytis cinerea*. **(A)** Relationship between mean fungal biomass (percentage of *B. cinerea* reads) and mean lesion area (mm²) at 72 hours post-inoculation (hpi) across ten host species. **(B-C)** Relationship between the total number of infection-responsive DEGs at 48 hpi and **(B)** mean lesion area at 72 hpi or **(C)** mean percentage of *B. cinerea* reads. **(D-E)** Relationship between the number of isolate-responsive DEGs at 48 hpi and **(D)** mean lesion area at 72 hpi or **(E)** mean percentage of *B. cinerea* reads. Infection-responsive DEGs represent the global response to infection independent of isolate identity, whereas isolate-responsive DEGs capture the variation driven by specific pathogen isolates. Solid lines represent linear regression fits based on Pearson correlation analysis, with shaded regions indicating 95% confidence intervals. Pearson correlation coefficients (*R*) and associated *p*-values are displayed to indicate the strength and significance of the observed associations.

**Figure S3:**
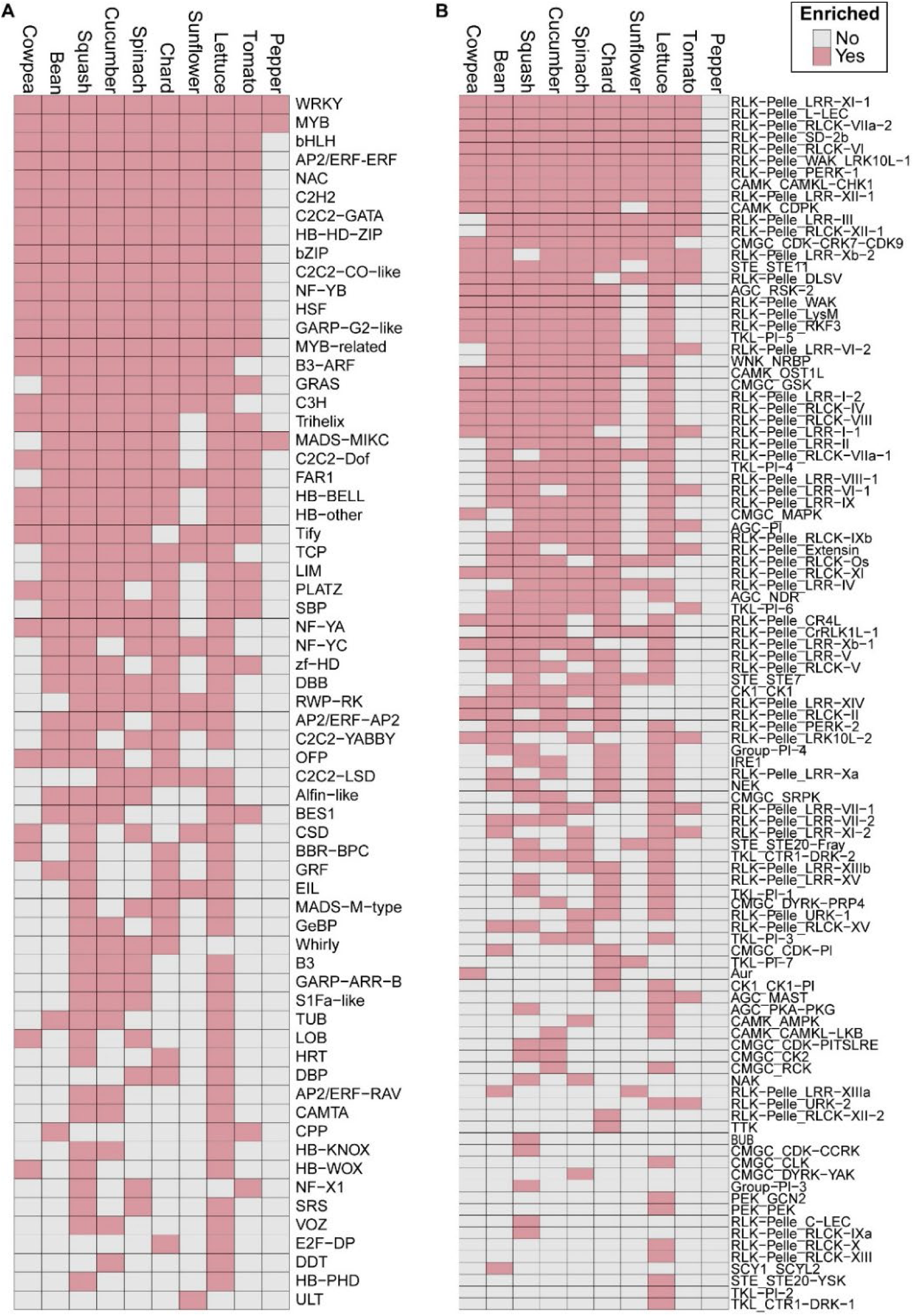
Enrichment of differentially expressed infection-responsive transcription factor (TF) families and protein kinases (PKs) across eudicots. **(A)** TF families and **(B)** PK enriched among infection-responsive differentially expressed genes (DEGs) in each host species. Rows correspond to families, ordered by the number of host species in which they are significantly enriched (from 10 to fewer). Columns represent the ten host species, arranged according to their phylogenetic relationships. TFs and PKs were annotated by mapping infection-responsive DEG sequences to the iTAK database (Zheng et al., 2016). Pink cells indicate significant enrichment (FDR ≤ 0.05), while grey cells indicate no enrichment.

**Figure S4:**
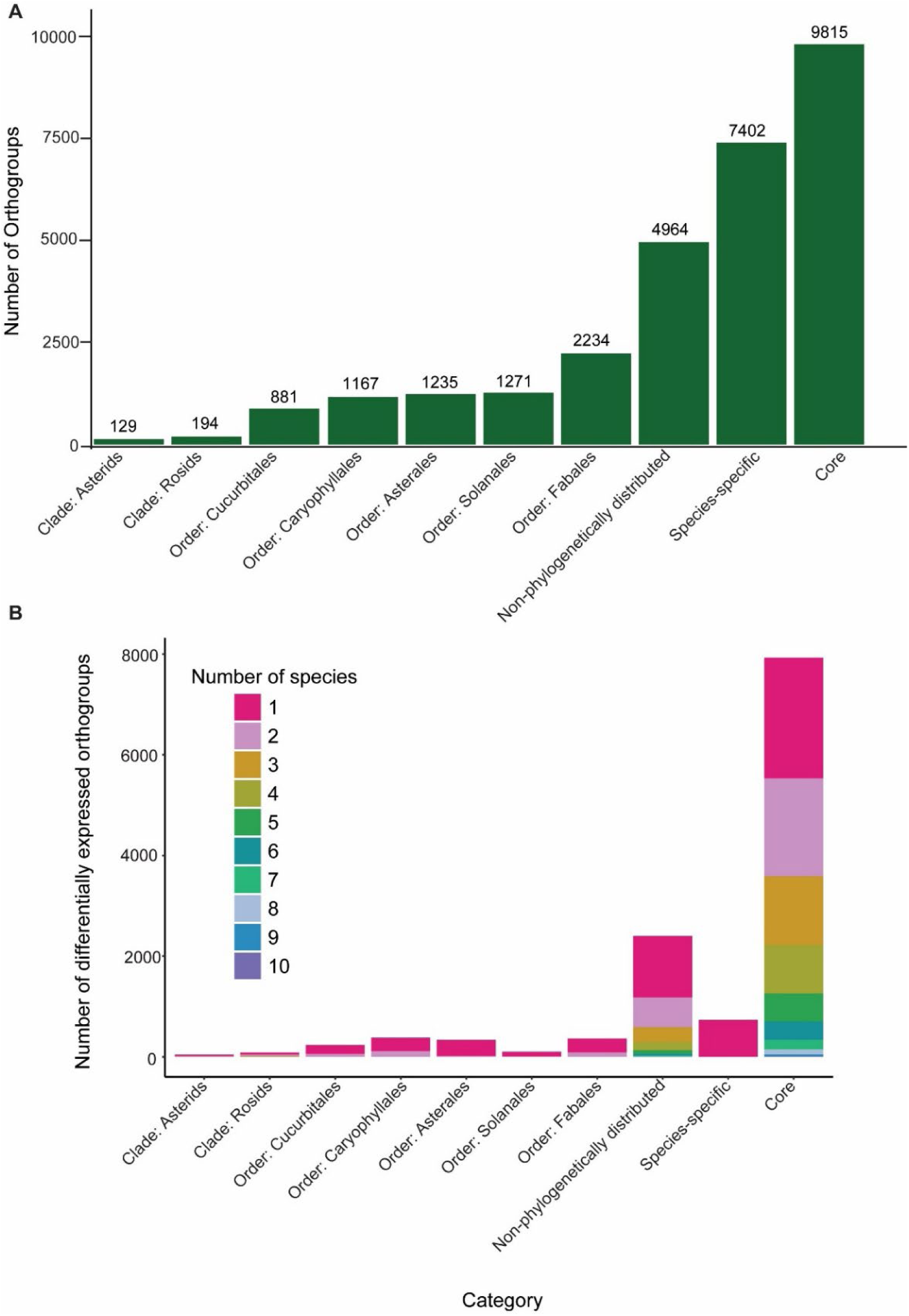
Orthogroup classification and differential expression across host species. **(A)** Classification of orthogroups identified by OrthoFinder across the ten eudicot host species at whole genome level. The bar chart categorizes orthogroups based on their phylogenetic distribution, showing counts for those conserved across all ten species, restricted to specific clades or orders, unique to a single species, or shared in other non-phylogenetic patterns. **(B)** For the categories defined in (A), the stacked bar plot shows the number of orthogroups that are differentially expressed in infection-responsive DEGs. The colour represents the number of species (from 1 to 10) in which a given orthogroup was found to be significantly differentially expressed.

**Figure S5:**
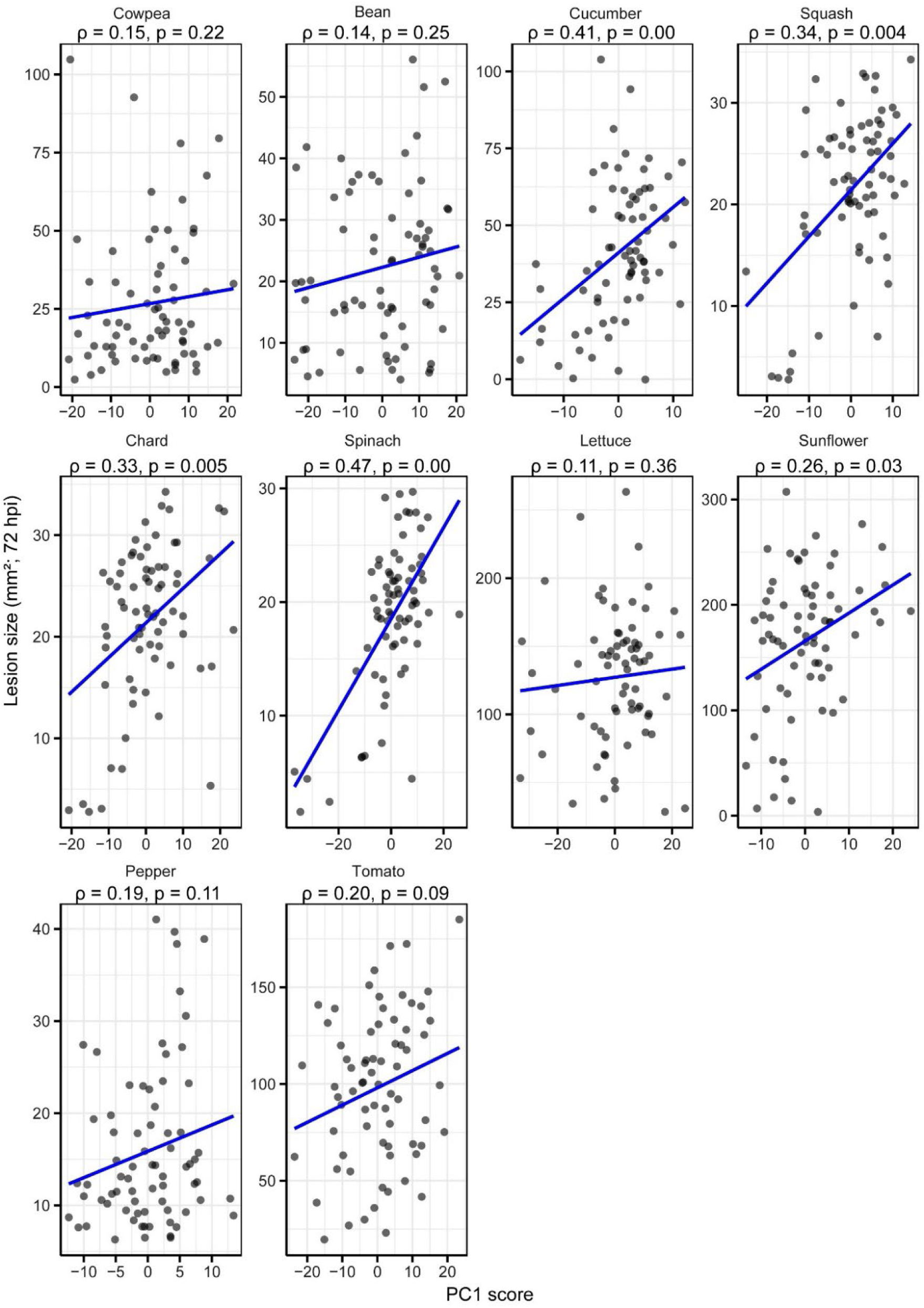
Correlation of PC1 scores with lesion size across host species. Scatterplots show the correlation between isolate-specific PC1 scores (from host transcriptional responses) and corresponding lesion sizes at 72 hpi within each host species. Each point represents a single isolate. Spearman’s correlation coefficient (ρ) and associated *p* value are indicated above each panel. Blue lines indicate linear regression fits for each host.

**Figure S6:**
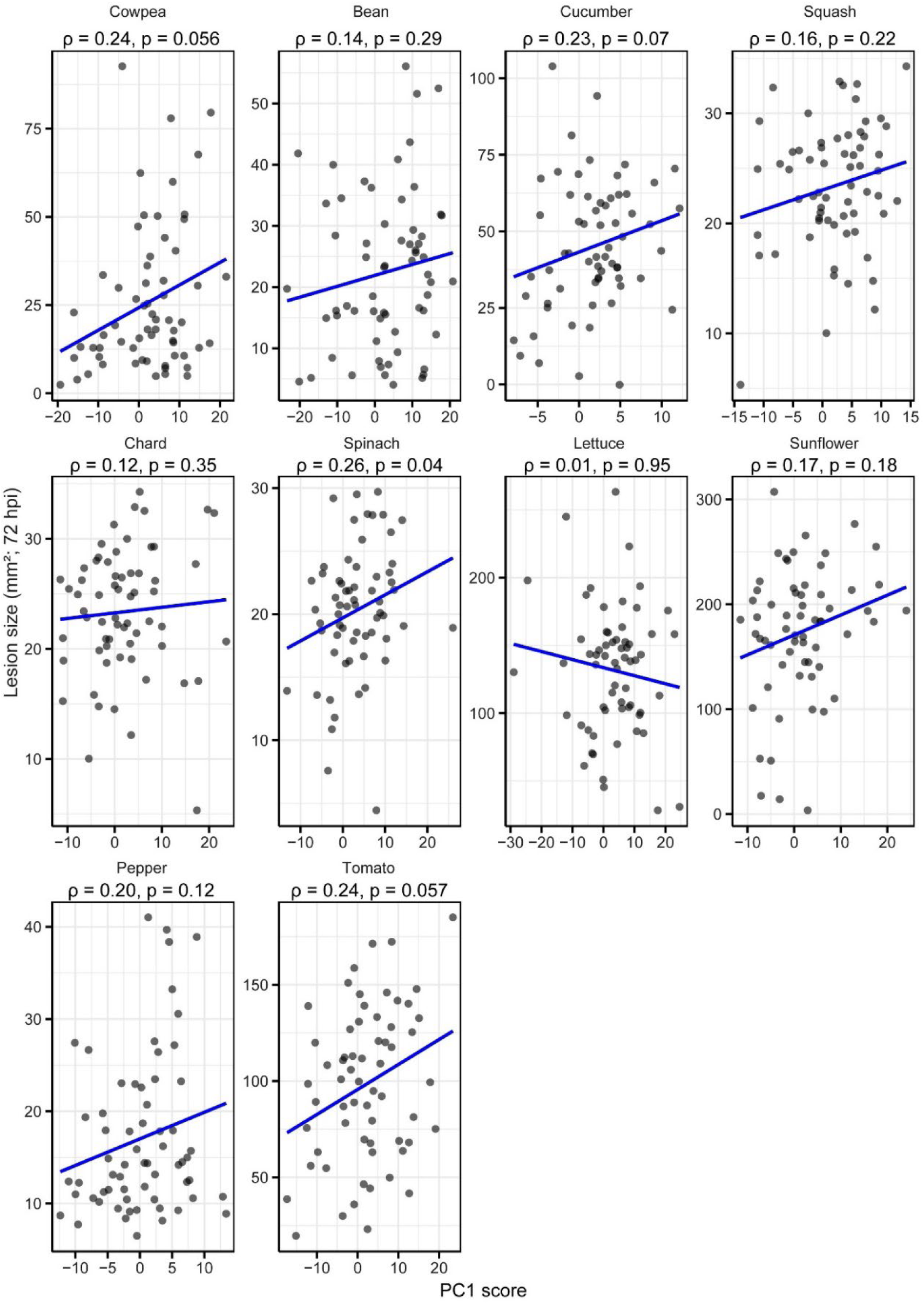
Correlation of PC1 scores with lesion size across host species after excluding weak-performing isolates. Scatterplots show the relationship between PC1 scores (from host transcriptional responses) and corresponding lesion sizes within each host species, after removing weak performing isolates (Gallo2, 01_02_18, 01_03_19, 01_04_21, FresaSD, 94_4, DavisNavel, UKRazz, 01_05_17) and Mock. Each point represents a single isolate. Spearman’s correlation coefficient (ρ) and associated *p* value are indicated above each panel. Blue lines indicate linear regression fits for each host.

**Figure S7:**
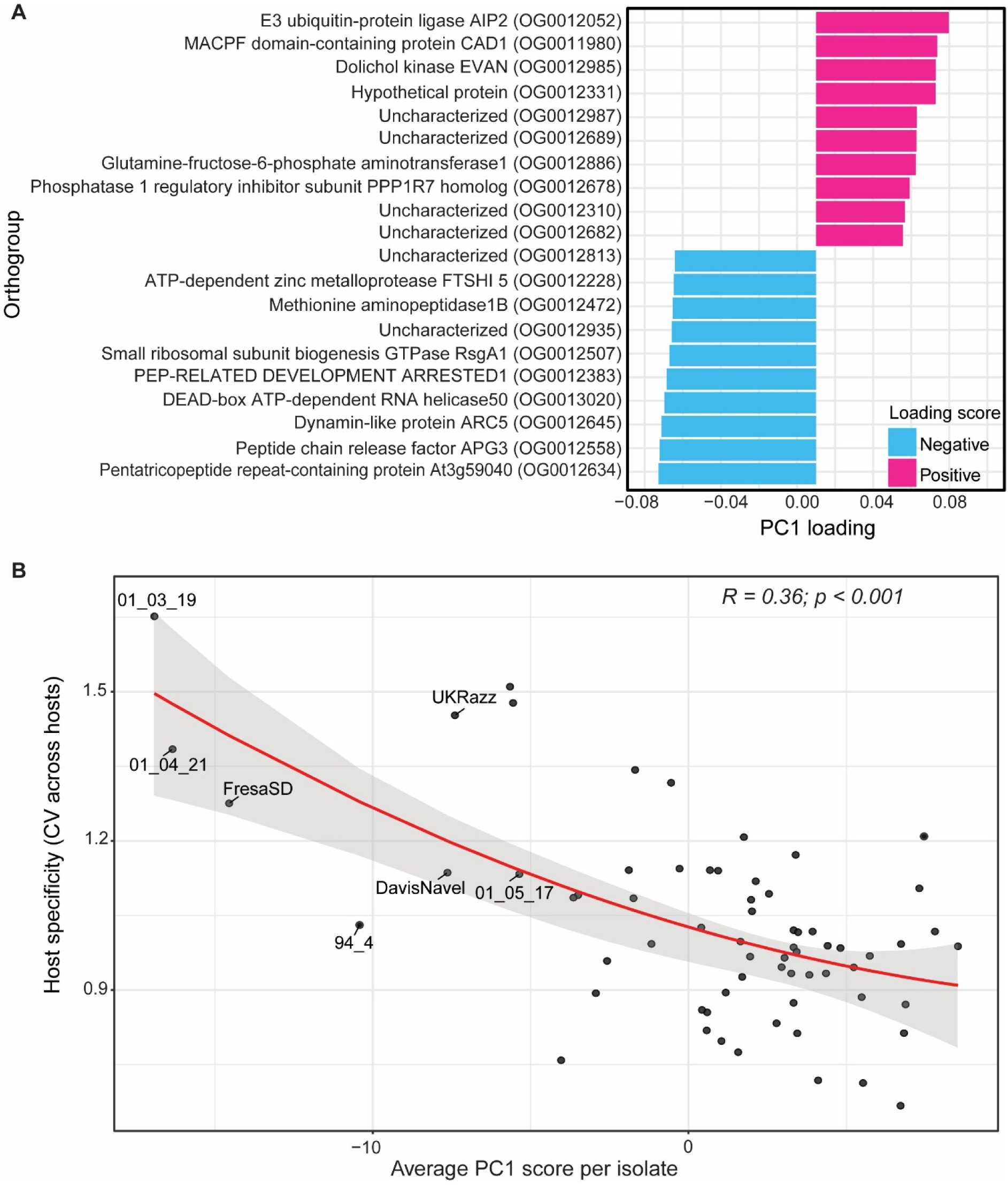
Principal component analysis (PCA) loading drivers and isolate-specific responses. **(A)** The top 10 positive and top 10 negative orthogroups contributing to Principal Component 1 (PC1) across hosts. Bars represent PC1 loading scores, with direction indicating positive (pink) or negative (blue) contribution. **(B)** Relationship between isolate average PC1 scores and host specificity, measured as the coefficient of variation (CV) of lesion size across hosts. The red curve shows the quadratic model fit, and the shaded area represents the 95% confidence interval. Dots represent individual isolates and weak-performing isolates are labelled for reference.

**Figure S8:**
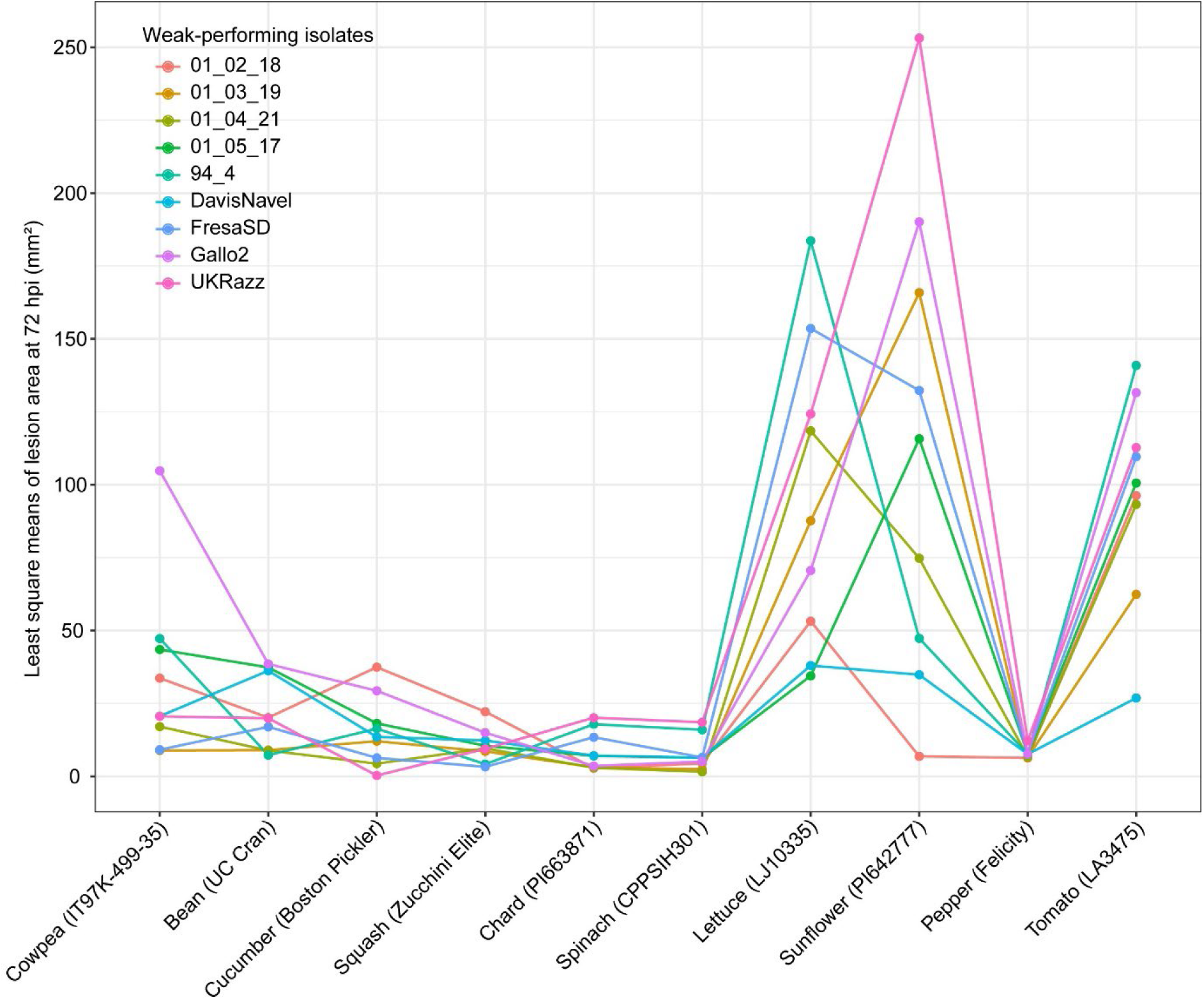
Lesion phenotypes of *B. cinerea* isolates eliciting weak host transcriptional responses at 72 hours post-inoculation (hpi). Line plot shows least squares mean lesion area in mm² at 72 hpi for *B. cinerea* isolates as having the lowest impact on host transcriptional response (based on PC1 scores). Each point represents lesion size measured on a single genotype per species, for which co-transcriptome was performed. Host species are shown on the x axis, with genotypes indicated in parentheses. Lines are coloured by isolate identity, as shown in the legend.

**Figure S9:**
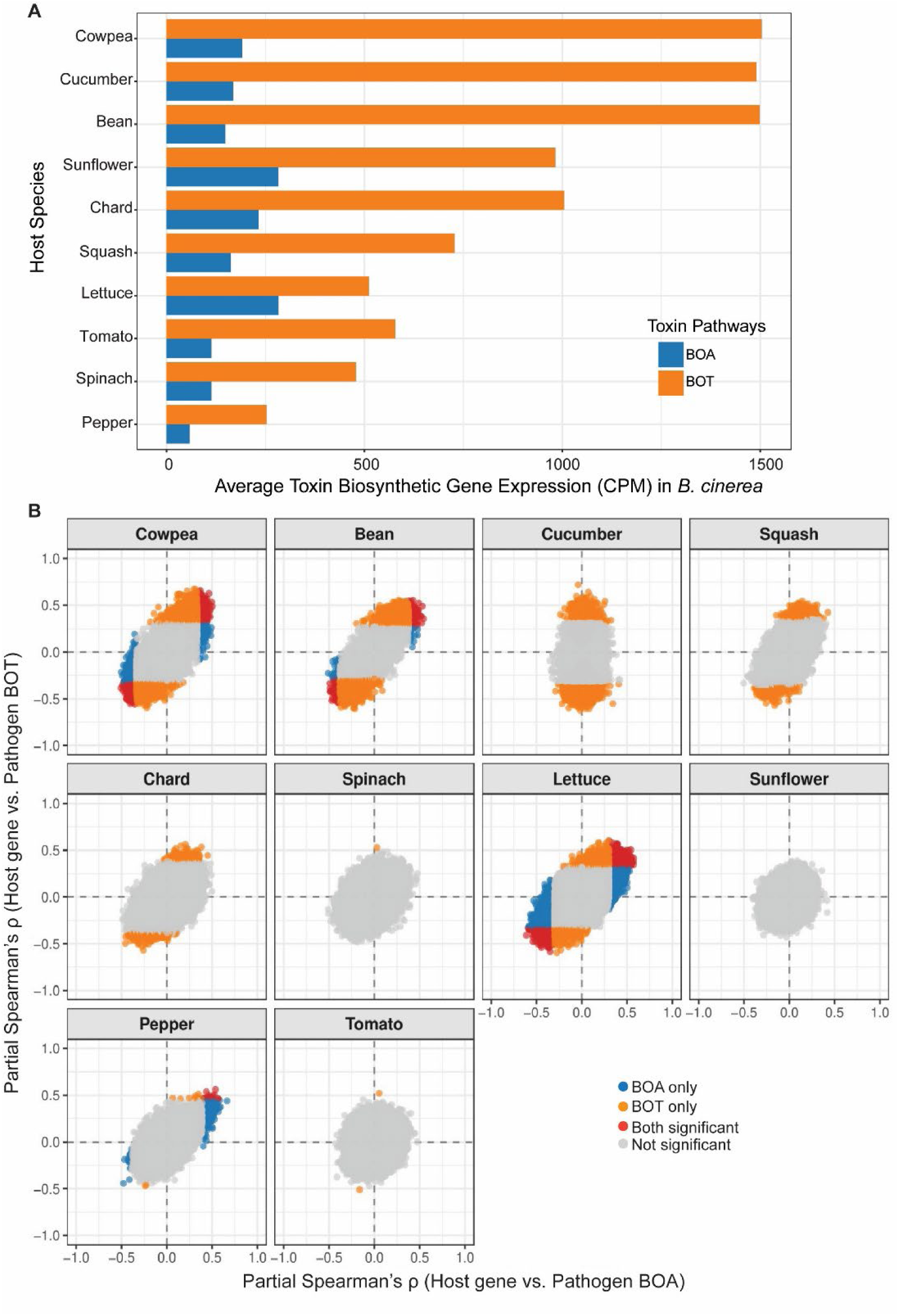
Pathogen phytotoxin biosynthetic gene expression and host responsive genes correlated with these phytotoxins. **(A)** Average transcript expression (CPM; counts per million) of the *B. cinerea* phytotoxin biosynthetic pathways i.e. BOA (*Bcin01g00010*-*Bcin01g00160*) and BOT (*Bcin12g06370*-*Bcin12g06410*) across host species. Blue bars represent BOA and orange bars BOT. For each isolate, expression was averaged across all genes within each biosynthetic cluster, and isolate-level averages were then averaged within each host species. This highlights substantial host-dependent differences in toxin pathway expression. **(B)** Spearman partial correlations (ρ) between host gene expression and pathogen toxin expression (BOA vs. BOT) for each host species, after correcting for general lesion associated genes (see material and method). Each dot represents one host gene. Genes significantly correlated (FDR ≤ 0.05) are colour coded as BOA only (blue), BOT only (orange), both significant (red), or non-significant genes (gray).

## Supplementary Tables

**Table S1:** List of reference genomes used for each host species as well as for *B. cinerea* in this study.

**Table S2:** Infection-responsive host differentially expressed genes (DEGs) and their phylogenetic classification across ten eudicot species. The table lists all host genes showing significant differential expression (FDR ≤ 0.05) in response to *B. cinerea* infection relative to mock controls at 48 hpi. Columns include the host species, gene and protein identifiers, assigned orthogroup IDs, and the phylogenetic classification of the orthogroup. Classifications indicate the evolutionary conservation of the gene family: Core (shared across all ten species), Clade-specific (restricted to Asterids or Rosids), Order-specific (restricted to a single order), Species-specific (unique to one species), or non-phylogenetically distributed (spanning multiple hosts without a clear phylogenetic pattern). Differential expression statistics (log2FC and adjusted p-value) were derived from comparisons between uninfected and infected samples.

**Table S3:** Summary of core infection-responsive orthogroups across eudicot hosts. The table details orthogroups identified via OrthoFinder that contain infection-responsive DEGs. Each row represents a single orthogroup, indicating the number of host species in which member genes are differentially expressed. Columns provide the counts of genes within each orthogroup classified by their transcriptional status during *B. cinerea* infection: downregulated (Down_inf-DEGs), not differentially expressed (Non_DEGs), or upregulated (Up_inf-DEGs). Functional annotations provide a putative biological role based on homology and gene descriptions.

**Table S4:** Summary of order-specific infection-responsive orthogroups. The table details orthogroups restricted to specific plant orders that contain infection-responsive genes. Each row represents a single orthogroup, indicating the number of host species within the order in which member genes are differentially expressed. Columns provide the counts of genes within each orthogroup classified by their transcriptional status during *B. cinerea* infection: downregulated (Down_inf-DEGs), not differentially expressed (Non_DEGs), or upregulated (Up_inf-DEGs). Functional annotations provide the putative biological role based on homology and gene descriptions.

**Table S5:** Host orthogroups correlated with *B. cinerea* toxin gene expression. The table lists host orthogroups containing genes whose expression levels were significantly correlated with the expression of fungal *Botrydial* (*BOT*) or *Botcinic acid* (*BOA*) biosynthetic gene clusters. Each row represents a single orthogroup. Columns include: Pattern (specifying the direction of correlation: Positive or Negative and the specific toxin involved); n_significant (number of host species exhibiting this correlation); significant species; n_sig_pos and n_sig_neg (counts of genes within orthogroup showing positive or negative correlations, respectively); Total_gene (total number of genes in the orthogroup); and Category/Subcategory (functional annotation and putative gene description).

**Table S6:** Information about genotypes of all eudicot species used for co-transcriptome. The germplasm ID corresponds to the germplasm collections (the U.S. Department of Agriculture Germplasm Resources Information Network [USDA GRIN], the Centre for Genetic resources in the Netherland [CGN], UC Davis Tomato Genetic Resource Centre [TGRC]), and other IDs represent IDs for species obtained from the commercial seed company. Information on the species and subspecies, common name, order, clade, and cultivar name is provided when available.

**Table S7:** Information for the 72 isolates of *B cinerea*. The name of the isolates, geographical origin with latitude and longitude coordinates (when available), the name and affiliation of the person who collected the isolates, the year of isolation and plant species (host) on which the isolate was collected are provided.

